# A rapid non-traditional approach for developing biologicals against drug-resistant bacteria and yeasts

**DOI:** 10.1101/2025.03.15.643420

**Authors:** Sanjiban K KBanerjee, Pratik Patil, Agastya Suresh Babu, Rija Nada, Gayatri Naik, Manjiri Shukla

**Author notes:** Corresponding author: Address for correspondence: Dr. Sanjiban K. Banerjee, AbGenics Life Sciences Pvt. Ltd. 4th Floor, Kant Helix, Bhoir Colony, Chinchwad, Pune 411033, India.

## Abstract

Development of effective, long-lasting antibiotics is a challenge due to the rate at which the pathogens acquire resistance, prompting the major pharma players to omit them from their portfolio. We present a non-traditional approach for rapidly developing anti-infective biologicals that is resilient to resistance. Antibody-drug conjugates were developed against three different pathogens *Pseudomonas aeruginosa* (ABG 14)*, Staphylococcus aureus* (ABG 16) and *Candida. albicans* (ABG 07) that were extremely specific and effective in neutralising them irrespective of their drug resistance profiles. Neutralising camelid antibody fragment (VHH) were isolated from immunised camel libraries by phage display that effectively neutralised the pathogens at a MIC 99 of 125 µg mL^-1^ *in vitro.* Antimicrobial peptides (AMP) were then conjugated to them by pathogen-specific cleavable linkers and the resultant molecules were 10 -20 times more effective due to a dual mode of action-the inhibitory action of the VHH on surface transporters and enzymes and the activity of the AMPs released by the pathogen surface proteases. Called AbTids, these molecules were extremely specific, stable in plasma and were activated in the presence of 10^4^ to 10^5^ CFU mL^-1^ of the pathogens and had an efficacy in the sub-micromolar range (6.25-12.5 µg mL^-1^, 250 nM) inhibiting their growth within 2 hrs. of administration. They were economically and efficiently produced in microbial production system as a single chain fusion protein. One of the molecules ABG 14 was characterised further and found to be pathogen specific with negligible frequency of resistance, was non-toxic, had the capability to destroy biofilms, and cleared a systemic infection in a mouse with a carbapenem resistant strain of *P. aeruginosa* at a dose of 5 mg kg^-1^. This strategy can be used to generate new hits against medically important pathogens in a matter of weeks by simply shuffling the components, using different VHHs and AMPs, bypassing years of expensive drug development efforts that can potentially rejuvenate drug discovery efforts against emerging superbugs.

## 1. INTRODUCTION

Drug resistant bacteria have been predicted to kill 10 million people, cause a loss of $100 USD trillion annually and wipe out the world’s GDP by 2-3.5% by the year 2050 (1). Recent reports predict a more serious threat, with 4.95 million deaths reported in 2019 alone. This was further exacerbated by the COVID-19 pandemic, which placed significant strain on healthcare systems and contributed to a rise in nosocomial viral and bacterial infections (2). Although the pandemic seems to be under control, the threat of drug-resistant pathogens remains as potent as ever. Among the main bacterial pathogens is a group known as the ESKAPE pathogens (*Enterococcus faecium, Staphylococcus aureus, Klebsiella pneumoniae, Acinetobacter baumannii, Pseudomonas aeruginosa, and Enterobacter species*) that are responsible for 1.27 million (2015 statistic) reported deaths, accounting for 73.4% of all fatalities attributed to AMR (3); the remaining due to other bacterial and fungal pathogens like Candida (4). Among the major pathogens in different geographies are methicillin resistant *S. aureus* which is responsible for as much as 54.9% deaths followed by *E. coli* with 30.4% and Candidiasis with 40%, but these figures vary from time to time. Another important pathogen *Pseudomonas aeruginosa,* is found in more than 50% of cystic fibrosis patients (5,6) and those resistant to carbapenems was responsible for increased morbidity and increased disease management costs. Multiple mechanisms of resistance like target mutations, antibiotic-inactivating enzymes, permeability barriers, active efflux of drugs might be operative in highly resistant strains (7), that can be transmitted across different gram-negative pathogens genetically as well (8). This has resulted in a rapid increase in the population of “superbugs” against the latest antibiotics, resulting in pathogens with more effective resistance mechanisms. Hence different types of intervention strategies - developing and deploying new drugs, vaccines, biologicals (9,10) are being implemented along with restricting and regulating the use of existing drugs via antibiotic stewardship programs (11) as a strategic approach to control their spread.

Traditional small-molecule antibiotics exert a selective force on the bacterial and fungal population that expedites the development of resistant strains within a few short generations, primarily driven by genetic mutations. Hence, multiple non-traditional approaches using biologicals (12) that are resilient to mutation-based resistance have been developed to overcome this limitation. Interest has been renewed in bacteriophages and microbiome engineering strategies as viable therapy options (13). Antimicrobial peptides have also been developed, particularly in combination with existing antimicrobials (14) along with antimicrobial antibodies (15). Though these strategies lack the mechanistic simplicity of antibiotics and are associated with some shortcomings, these approaches can be improved to enhance their druggable properties and mode of action which will hold out against simple point-mutation based resistance mechanisms. This would result in antimicrobials with longer and effective product lifecycle similar to that of vaccines that are effective even decades after introduction as compared to antibiotics that become resistant in few short years in the clinic (16).

We have developed a new class of antibody-drug conjugates called ‘AbTids’ using camelid variable heavy chain (VHH) antibody fragments, combined with cleavable Antimicrobial Peptides (AMP) and tested this concept for efficacy and specificity against three important microbes in the CDC and WHO critical list. Using different pathogen-specific VHH and the AMP histatin, we demonstrate a strategy for designing precision antimicrobial biologicals against multiple drug-resistant pathogens using a common design approach. The VHH and AMP have been joined together by a pathogen specific protease-cleavable linker that is activated specifically on contact with the pathogen resulting in the release of the bacteriolytic AMP from the conjugate that neutralises the pathogen rapidly.

Camelids (camels, llamas, and alpacas) produce unconventional single heavy chain antibodies that are devoid of a light chain (17,18). The antigen binding part-VHH is extremely small with a size of 15 kDa, has a high affinity for antigens and ability to bind to cryptic epitopes (19). They have higher thermal and chemical stability and can be economically produced by simple microbial systems that have enabled their use for therapeutic and diagnostic applications (20). Their small size, stability, and low immunogenicity makes them suitable for therapeutic applications like neutralising antibodies and can be administered as antibody-drug conjugates (21). They have been used for *in vitro* diagnostic assays and for *in vivo* applications like whole body imaging in animals by attaching them to radionuclides (22). A humanised bivalent VHH, Cablivi is in the market as a therapeutic molecule for treating clotting disorders (23) and thirteen molecules are in various stages of preclinical and clinical development attesting to their usefulness as a new class of therapeutic molecules (12).

Multicellular organisms use antimicrobial peptides as a first line of defense due to their broad killing activity against fungi, bacteria, yeasts, cancer cells and even viruses (22). They are extremely toxic at higher concentrations and some like colistin are used as a last resort drug for control of superbugs in critical care settings (24). Another AMP, the salivary histatin, used in this study, is extremely potent against a wide spectrum of bacterial and fungal pathogens but has not been used for therapeutic use (25). Found in the saliva of humans and some higher primates, histatins target the mitochondria and initiate cell death by inducing the non-lytic loss of ATP leading to disruption of cell cycle (26). Multiple modes of action of histatins are seen in different pathogens, with membrane disruption in *P. aeruginosa* and targeting of mitochondria (27), inhibiting respiration, accelerated formation of reactive oxygen species (28) to the release of intracellular potassium in *Candida* (29). They have been demonstrated to be effective against all the ESKAPE pathogens attesting to their usefulness as broad-spectrum antimicrobials (30).

In this study we demonstrate the effectiveness of AbTids against three different pathogens - Gram positive *Staphylococcus aureus*, Gram negative *Pseudomonas aeruginosa,* and yeasts *Candida albicans* and *Candida auris sp*. - keeping the AMP constant and varying the VHH and cleavage sites depending on the chosen pathogens and comparing the efficacy and specificity of neutralisation. AbTids were found to be extremely specific, with an efficacy in the sub-micromolar range, showing dose dependent antimicrobial activity that are non-toxic to human cells and are plasma stable. One of the AbTids ABG 14 was characterised further and found to destroy biofilms, resilient to resistance and cleared systemic infection of a carbapenem resistant *P. aeruginosa* from a neutropenic mouse with significant reduction of bacterial counts within 24 hr. of administration establishing the effectiveness of the design concept of AbTids that can be used to develop other anti-infective biologicals rapidly against emerging threats.

## 2. MATERIALS AND METHODS

### 2.1. Bacterial strains

Pathogenic bacteria *P. aeruginosa*, *S. aureus* and yeast *Candida sp*. were purchased from ATCC. Locally sourced bacteria were deposited to National Centre for Microbial Resources, Pune and were assigned MCC numbers. Probiotic strains were purchased from Microbial Type Culture Collection (India). Multidrug resistant *P. aeruginosa* (BAA 2108) was purchased from ATCC, Carbapenem resistant *P. aeruginosa* (strain MCC 50428), was isolated from patient sample and deposited in National Centre for Microbial Resources, Pune, India, and *P. aeruginosa* (Schroeter) Migula (ATCC 27853) was used as a model strain for drug sensitivity studies. Methicillin resistant *S. aureus* MRSA (strain MCC 50811), and methicillin sensitive MSSA (MCC 51005) were isolated from patient samples. *C. albicans* (MTCC 227) was purchased from microbial type culture collection (India) and *C. auris* was purchased from ATCC (MYA 5002). The probiotic bacteria were purchased from Microbial Type Culture Collection (MTCC) India *L. plantarum* (MTCC 1325), *L. casei* (MTCC 12468), *L. acidophilus* (MTCC 10307), *L. fermentum* (MTCC 5898), *E coli* (MTCC1610 and *B. subtilis* (MTCC 2389). For cloning and expression of AbTids and VHH, *E. coli* ER 2738 (NEB, UK), *E. coli* BL21 (DE3), *E. coli* C 41, helper phage (M13 KO7) was purchased from vendors and growth media and antibiotics procured from Hi Media (India).

### 2.2. Generation of Camelid antibody libraries and isolation of pathogen-specific hits

Three Indian dromedary camels (*Camelus dromedarius*) were infected with an inactivated meropenem-resistant *P. aeruginosa* MCC 50428, methicillin resistant *S. aureus* MCC 5081, and *C. albicans* (MTCC 227) and three immunised libraries (CIL) CIL-Pa for *Pseudomonas aeruginosa*, CIL-Sa for *Staphylococcus aureus* and CIL-Ca for *Candida albicans* were generated using the protocol described earlier (53). Ten randomly selected colonies were subjected to colony PCR to check the presence of the VHH insert and digested with *BstN*1 to determine the diversity that was found to be in the 90% range (Fig. not shown). The libraries were then stored at -80°C in LB + 50% glycerol till further use.

Enrichment of phages expressing the hits was carried in immunotubes (Thermo Fisher Scientific, USA) pre-coated with whole-cell *P. aeruginosa, S. aureus* and *C. albicans.* Three consecutive rounds of bio-panning were done for each of the three libraries. Eighty colonies were randomly picked up from the enriched libraries and screened for antigen binding by phage-ELISA using anti-M13-HRP conjugated antibodies (Genescript, USA) (31). Recombinantly purified targets were coated on an immune plate and bound phages detected by anti-M13-HRP conjugated antibody (working concentration 0.4 µg mL^-1^). Wells giving 4-times more readings than the blanks were identified as hits and used for expression and analysis.

### 2.3. Expression and purification of target antigens

For expression and purification of target antigens against the three pathogens, primers were prepared against a component of C4-dicarboxylate transporter gene of *P. aeruginosa* and enolase genes of *C. albicans* and *S. aureus*. The full-length genes were extracted by PCR, cloned and expressed in pET28c vector in *E. coli* C 41 (D3) and the histidine tagged targets were purified using metal affinity chromatography. The protein bands of C4-dicarboxylate transporter (33 kDa) and both enolases (47-48 kDa) were run on SDS-PAGE, and extracted by electro-eluter (Model 422 Bio-Rad, USA).

### 2.4. Design of AbTids against three pathogens

All the three AbTids had the same basic design consisting of three components. The sequences of all the pathogen specific VHH hits were examined for the completeness by locating the start and stop codons as well as by demarcating the framework regions. (Framework 1: D V Q L Q E S G G D S V Q E G D S L T L S C K V S, Framework 2: C M A W F H Q A Q G K E N E A V A A, Framework 3 : T T Y A E S A K N R F T I A R D K L K S V V T L Q M N S L K V E N T G L F R C A A and Framework 4: Q G Q G T Q V T V S G). VHH fragment (Ca C11h for *C. albicans*, Ps C23 for *P. aeruginosa* and Sa C19 for *S. aureus*) against a surface antigen of individual pathogens were linked to a pathogen-specific protease cleavage site followed by the antimicrobial peptide histatin as indicated in the Fig. 1c. To give more flexibility to the design, flexible amino acids glycine and serine were used in two forms – a long linker (G4S)3 and a short linker G4S were used with the protease cleavage site. Two forms of histatin, an intact fragment and the active component were used (32,33) and the gene constructs were synthesized from Genescript (USA), cloned and expressed in *E coli*. Three AbTids were developed for testing – ABG 07 for *Candida sp.*, ABG 14 for *P. aeruginosa* and ABG 16 for *S. aureus*.

### 2.5. Expression and purification of the AbTids in *E coli*

2.5.1. Expression and isolation of inclusion bodies

The pET28c vector containing the three AbTid genes ABG 07, ABG 14 and ABG 16, were transformed in *E. coli* C41 (D3) cells. A single transformed colony for each was induced with 1 mM isopropyl β-D-1-thiogalactopyranoside (IPTG) for 5 hr, the culture was harvested and suspended cells were disrupted by sonication on ice for 20 min. The pellet was suspended in urea wash buffer consisting of 2 M urea, 20 mM Tris-HCl, 0.5 M NaCl, 2% Triton™ X-100, 1X protease inhibitor cocktail, pH 8.0 and sonicated again for 20 min. with cycle of 5s pulse and 15s rest and inclusion bodies of the AbTids were collected by centrifugation at 12000 x g for 10 min at 4°C. Solublisation and refolding of inclusion bodies The pellet (unpurified inclusion bodies of AbTid) was suspended in solublisation buffer (20 mM Tris-HCl, 6 M Guanidine-HCl, 0.5 M NaCl, 5 mM BME, pH 8.0) and supernatant was collected after centrifugation at 12000 x g for 30 min. at 4°C and loaded on Ni-Sepharose high affinity column (His-Trap TM) and protein elution was carried out with linear gradient of imidazole with concentration ranging from 0 M -0.5 M in elution buffer (20 mM Tris–Cl, 0.5 M NaCl, 0.5 M imidazole, pH 8.0). The fractions containing the peaks were collected and imidazole was removed by 10K centrifugal filter (Vivaspin® 20 10 kDa MWCO, Cytiva Sweden) using buffer containing 20 mM Tris-Cl, 0.1 M NaCl pH 8.0 and the eluates were further purified by size exclusion chromatography (Superdex-S75). The purified inclusion bodies were refolded by drop-wise dilution method in cold (4℃) refolding buffer containing 25 mM Tris, 0.1 M Sodium chloride, 0.5 M L-arginine, 4 mM GSH (Glutathione), 0.4 mM GSSG (Glutathione disulfide), pH 7.2 and incubated for 24 hr. at 4°C. The concentrated refolded AbTids (active form) was analysed on 15% SDS-PAGE. The endotoxin concentrations were measured with LAL endotoxin quantitation kit (Thermo Fisher Scientific, USA). The final concentration of AbTids were measured by micro-bicinchoninic acid assay and stored in 50 mM phosphate-citrate buffer containing 50 mM phosphate-citrate, 0.1 M NaCl, 10% glycerol, pH 7.2 at -20° C for further use.

### 2.6 Characterisation of AbTids

#### 2.6.1 Binding and specificity assays

300 ng of recombinantly produced Candida enolase, Staphylococcus enolase and Pseudomonas C4-dicarboxylate transporter were coated in triplicates on ELISA plates along with negative control containing no antigen, blocked (with 3 % skim milk, incubated in 37℃ for 1-hr), followed by incubation with AbTids ABG 07, ABG 14 and ABG 16, washed thrice in PBST followed by incubation with secondary mouse anti-VHH HRP conjugate (1:3000) (Genescript, USA) and colour developed after washing of unbound secondary antibody by addition of the substrate (1:9) 3,3′,5,5′-Tetramethylbenzidine (Sigma-Aldrich, USA) for 10-15 minutes. The reaction was stopped and absorbance at 450 nm was measured in a microplate spectrophotometer (Gene 5, BioTek Epoch).

#### 2.6.2 Detection of AbTid activation by cleavage assays

For each of the pathogens, a single colony was inoculated in 2 ml of Mueller Hinton media (MH broth) with added FBS, grown overnight at 37°C till OD=1 at 600 nm (10⁹ CFU mL^-1^) and supernatant containing the proteases are collected by centrifugation at 9000 x g and filtered with 0.2-micron filter to remove the pathogens. 500 µl of supernatant and 15 µg (in 15 µl) of the AbTids were incubated for 2, 4 and 6 hours at 37°C. In the second set, the culture was serially diluted to achieve increasing CFU mL^-1^ of pathogens (from 10² to 10⁹) and supernatants and the AbTids were incubated for 2-hours at 37°C and 20 µL of each CFU mL^-1^ (10²,10³,10⁴,10⁵,10⁶,10⁷,10⁸,10⁹). Samples were then loaded on a 15% SDS-PAGE, analyzed for integrity of the intact molecules via band-size comparisons.

#### 2.6.3 Plasma stability and toxicity to human cells

Plasma stability assay was performed on the AbTids using complement inactivated human plasma that was incubated at 37°C with same amounts of the three AbTids. 4.76 µg in 100 µL (4.76 µM) of the three AbTids were added and analysed at 0, 2, 4, and 24 hr. respectively on 15% SDS PAGE (Bio-Rad Laboratories, Inc. USA) to check for the integrity by observing for fragmentation or cleavage-based activation and change in size of the molecules.

The toxicity of one of the AbTids, ABG 14 was tested on a monolayer of HEK 293T cells after incubating it with three different concentrations (MIC dose, 5X MIC and 10X MIC dose) of the AbTid 12.5 µg mL^-1^, 62.5 µgmL^-1^ and 125 µg mL^-1^, followed by MTT assay after 24 hr. of incubation. 1% Triton X-100 was used as a positive control.

#### 2.6.4. Biofilm assay

The biofilm formation assay was performed by following a protocol mentioned earlier with slight modifications (34). Overnight cultures of carbapenem resistant *P. aeruginosa* (MCC 50428) were diluted in LB to OD_600_ of 0.02. 100 µL of the sample suspensions were mixed with 25 µg mL^-1^ of ABG 14 and applied into 96-well flat-bottom microtiter polystyrene plates (Becton, Dickinson) and incubated for 16 hrs at 30^◦^C. The next day, they were centrifuged (3000 rpm, 10 min, washed with PBS three times and biofilms were visualized by staining with 125 µL of 0.1% crystal violet for 15 min. at room temperature. The amount of biofilm was quantified after adding 100 µL of 95% ethanol to dissolve it and measured at OD_570_. A bacterial sample treated with BSA was used as a negative control.

#### 2.6.5. Microbiological assay of efficacy and specific activity of AbTids

The freshly grown cultures of OD_600_ 1 of *P. aeruginosa* BAA 2108 (Multidrug resistant), *P. aeruginosa* 27853, *C. albicans* (MTCC 227), *C. auris* (ATCC MYA 5002), Methicillin resistant *S. aureus* (MCC 50811) and Methicillin sensitive *S. aureus* MCC 51005 in Mueller Hinton (MH) broth were serially diluted up to 50 K cells mL^-1^. 100 µL of each of the pathogens were added to MH broth with 25 µg mL^-1^ of each of the AbTids, incubated at 37°C for 16 hr. and the growth quantitated by OD_600_ and dilution plating. All the three AbTids were added to each of the three pathogens to check for the specificity of action. For time-kill assays ABG 14 (6.25µg/ml), PsC23 (25 µg/ml) and meropenem (8µg/ml) were added to for both the *P. aeruginosa* strains. (1 x 10^6^ CFUmL^-1^) and growth monitored for 24 hrs. At time intervals 0, 2, 4, 8, and 24 h, 10 µL samples were withdrawn from each well and plated on MH agar to determine the CFU mL^-1^. To determine if the AbTids had inhibitory effect on non-pathogenic strains, the activity of the Pseudomonas AbTid ABG 14 was checked on the *P. aeruginosa* strains BAA 2108 and ATCC 27853 and the probiotic bacteria normally present in the gut-*L. casei, L. fermentum, L. acidophilus* and *L plantarum* as well as two unrelated bacteria *E. coli* and *B. subtilis.* All the bacteria (OD_600_ 0.1) were grown in the presence of 12.5 µg mL^-1^ of ABG 14, and growth quantitated by measuring the OD_600_ after 16 hr.

#### 2.6.6. Adaptive resistance studies of *P.aeruginosa* ATCC 27853 and MCC 50428 to AbTid ABG 14 and meropenem

Carbapenem sensitive *P. aeruginosa* ATCC 27853 and resistant MCC 50428 were serially passaged at a fixed sublethal concentration of the two molecules to allow them to adapt. The bacteria were grown overnight, OD_600_ was adjusted to 0.2, meropenem and ABG 14 were added in concentrations of 0.5 µg mL^-1^ and 3.125 µg mL^-1^ respectively, incubated overnight in MH broth and then serially passaged for 9 generations and OD_600_ plotted after every 3 generations (G3, G6 and G9) along with dilution plating to determine the growth characteristics in persistent doses of the drugs.

#### 2.6.7. *In vivo* efficacy studies

Animal experiments were carried in collaboration with PRADO Preclinical and Research Facility, according to the guidelines and approval of the Institutional Animal Ethics Committee (Approval number IAEC-19-015). Pathogen-free 6-8-weeks-old BALB/c mice (female) were housed in polypropylene cages and maintained under controlled conditions (23°C – 25°C, 50 – 55% humidity and 12 hr. dark and light cycles). They were fed with chow diet and water *ad libitum*. Before infection, neutropenia was induced in these mice by injecting two intra-peritoneal doses of cyclophosphamide on day-4 (150 mg kg^-1^) and day-1 (100 mg kg^-1^). On the day of infection, an overnight grown culture of meropenem-resistant *P. aeruginosa* (MCC 50428), 0.5 McFarland standard suspension (equivalent to a density of 10^8^ CFU mL^-1^) was prepared in a sterile lactate ringer’s solution. Infection was induced by injecting 50 µL bacterial suspension into tail vein of mice using 27-guage needle.

A total of five groups (*n* = 4) of mice were used. One-hour post-infection, mice received the 5 mg kg^-1^ dose of ABG 14 BID and their condition was monitored for 72 hr. as described in Table 1 and survival of the control and treated animals was monitored. Heparinised blood samples were collected aseptically after 24 hr., centrifuged at 2,000 x g for 5 min. and a suitable dilution was plated on MH agar, and incubated at 37^0^C, 24 hr. for determination of the bacterial load (CFUmL^-1^). The 5^th^ group of mice was administered purified ABG 14 (equivalent to 5 mg kg^-1^ of body weight) without inducing any infection and examined for the preliminary signs of toxicity (data not shown).

**Table 1:** Percentage homology of the different pathogen surface targets used in the study.

| Organism | Homology with Staphylococcus enolase (%) | Homology with Candida enolase (%) | Homology with Pseudomonas enolase (%) | Homology with human enolase (%) |
| --- | --- | --- | --- | --- |
| *S. aureus* | 100 | 52.90 | 62.35 | 49.65 |
| *C. albicans* | 52.90 | 100 | 52.35 | 63.59 |
| *P. aeruginosa* | 62.35 | 52.35 | 100 | 53.32 |
| *H. sapiens* | 49.65 | 63.59 | 53.32 | 100 |

## 3. Statistical analyses

The statistical analysis of data was done using GraphPad prism (v 9.5) and (one way) <u>an</u>alysis <u>o</u>f <u>va</u>riance (ANOVA) was applied. Statistical significance threshold was set at *P* < 0.05 and confidence level set was 95%.

## 3. RESULTS

### 3.1 Design considerations and components of the VHH-AMP conjugates (AbTids)

The two major components of the AbTids, the VHH and the AMP were linked by a protease cleavable linker) so that the peptide is released by the pathogen proteases resulting in an effective bactericidal action (Fig. 1a). Specificity was ensured by carefully choosing the protease cleavage sites that are cleaved only in the presence of the target bacteria but are stable in human plasma. Dual checkpoints of pathogen specificity by antibody binding and cleavage by pathogen protease ensured a precisely targeted lethal dose. After attachment of the AbTid to the pathogen outer membrane pathogen killing was effected by the neutralising antibody and by the bactericidal action of the released AMP. Depending on the properties of attached AMP, bacteriolytic action could be due to cell wall lysis, biofilm destruction, interference with DNA functions. Due to this dual mode of action, it would be difficult for the pathogen to mount a simple point mutation-based resistance mechanism as with small molecule antibiotics (Fig. 1b). AbTids can thus potentially be resilient to resistance as they do not penetrate the cell, thereby negating the effect of permeability barrier and efflux mediated resistance as well. They can be assembled synthetically, produced in microbial production systems, and tested in a matter of weeks, considerably shortening the drug discovery process. Multiple emerging drug-resistant pathogens can be targeted as well, by simply changing the antibodies and the cleavage sites, ensuring a robust pipeline that can outpace emerging superbugs. The design principle and logic for using the different components for each of the AbTids are enumerated below.

**Fig.1a.**
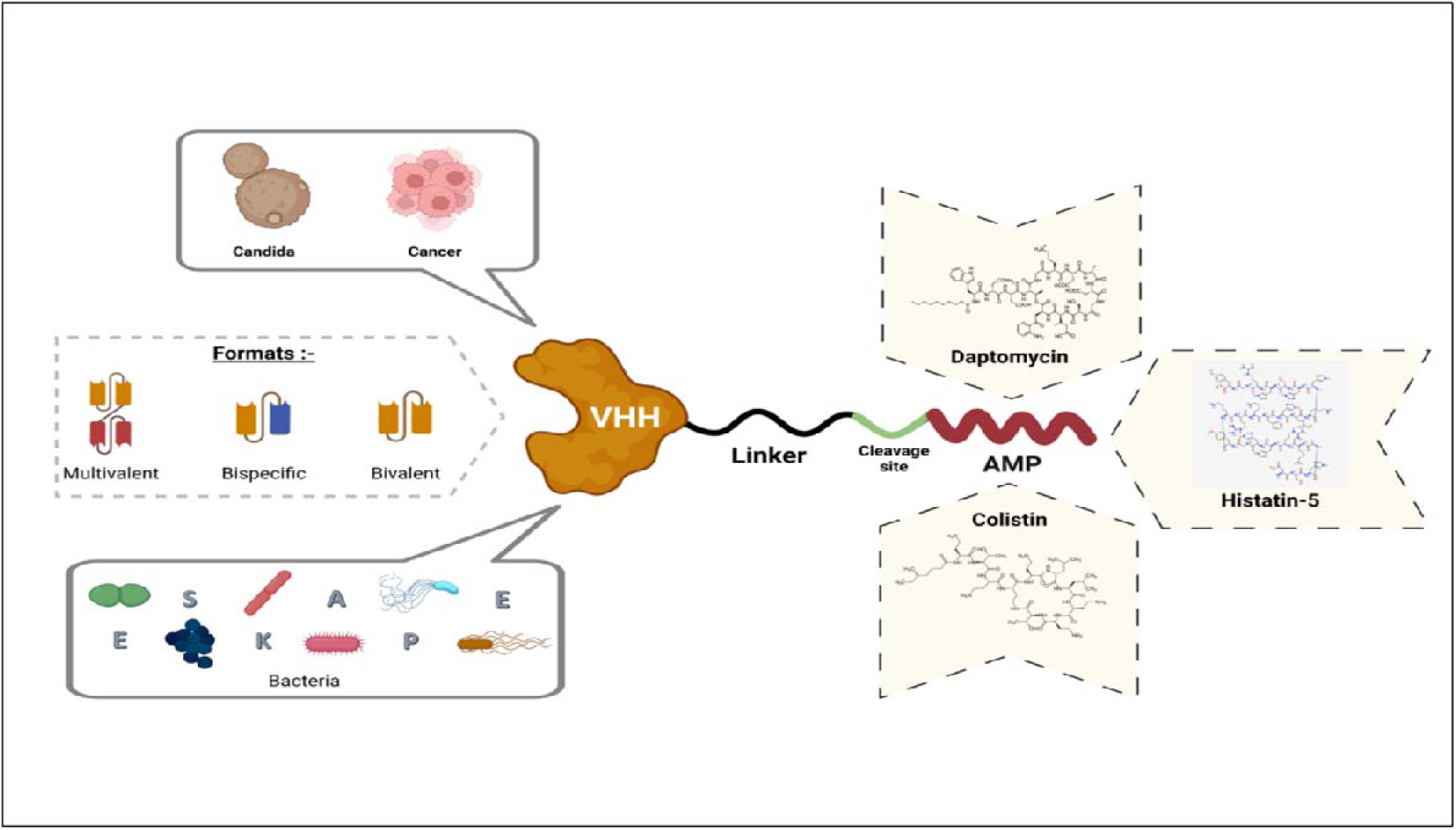
Design and construction of the VHH-AMP conjugates (AbTids). The VHH is camelid antibody fragment against bacterial pathogens like ESKAPE, fungal or cancer cells. The VHH linked by a pathogen-specific protease cleavable linker to the antimicrobial peptide (AMP) either by chemical conjugation or produced as a fusion protein if the peptide is linear. A single VHH can be used or bivalent or bispecific versions can be made by fusing two fragments by a flexible linker. AbTids can be produced in microbial production systems, is modular and the three components can be shuffled to generate variants to target multiple pathogens irrespective of their drug resistance profiles.

**Fig.1b.**
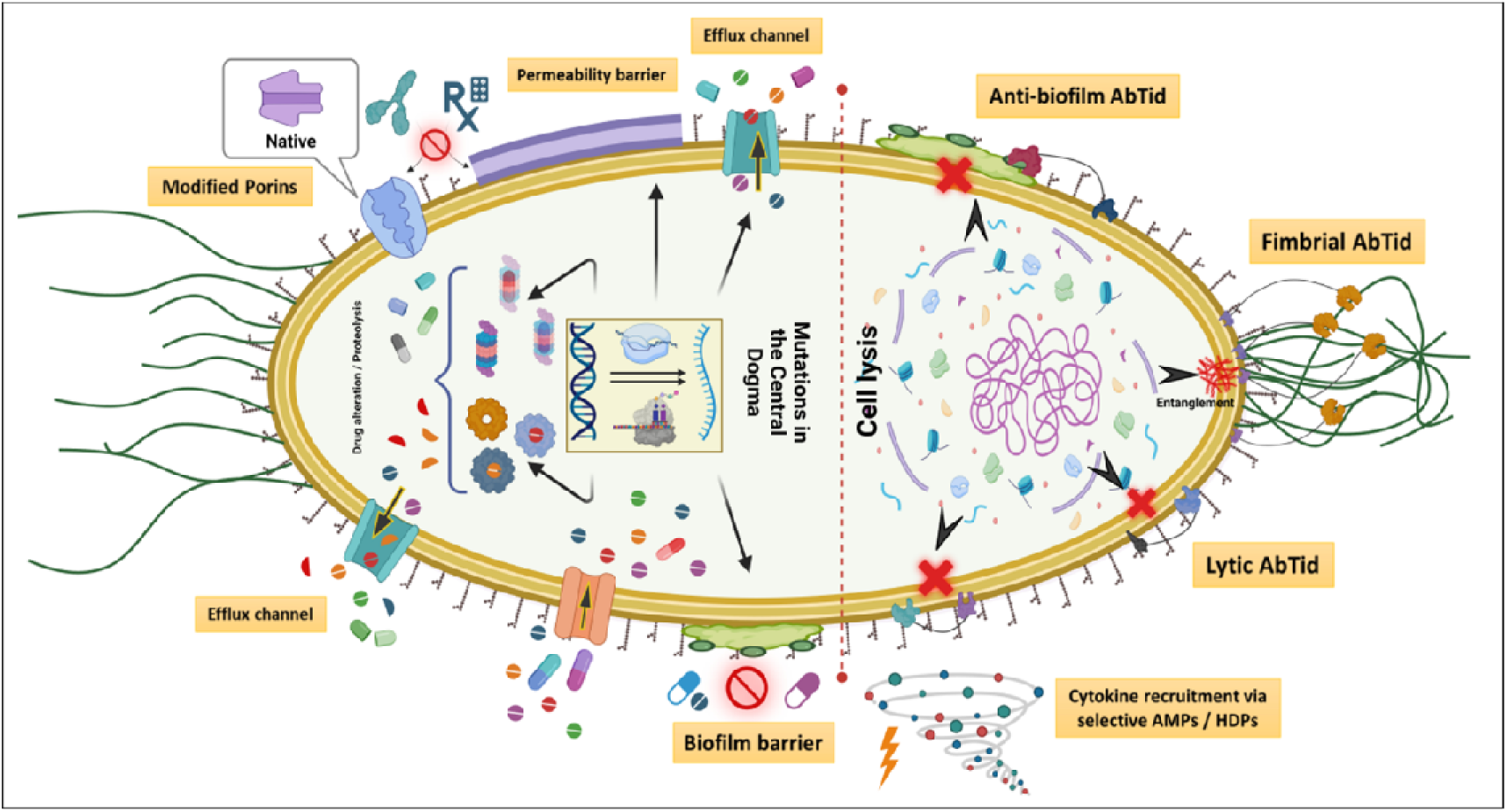
Non-traditional antibiotic properties of AbTids. Resistance mechanisms in bacterial pathogens to traditional small molecule antibiotics (left) and the different modes of action of the conjugates (right) making them resilient to resistance. AbTids do not enter the cell, have multiple attachment points thereby resisting simple point mutation-based resistance mechanisms and combine different antibacterial actions for effective pathogen neutralisation.

**Fig.1c.**
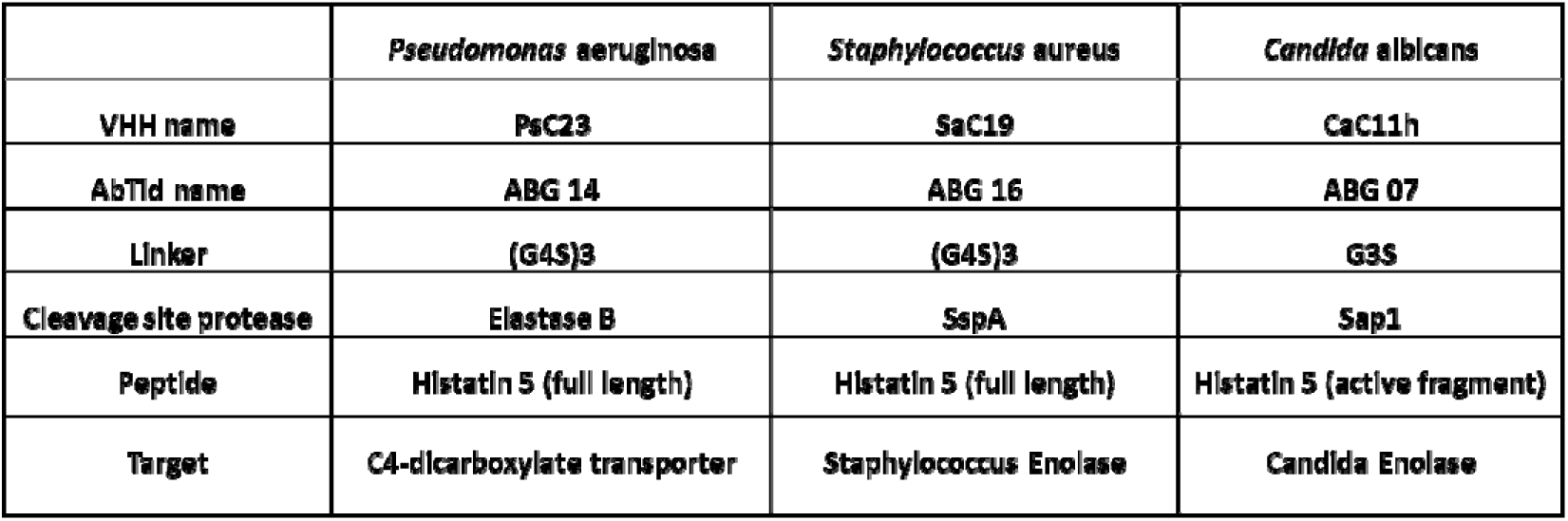
Components of AbTids for *P. aeruginosa* (Gram negative), *S. aureus* (Gram positive) and *C. albicans* (fungus). The three pathogens were chosen to demonstrate the robustness of the common design principles for all the three molecules. The VHHs are joined to the antimicrobial peptide histatin by a linker containing a pathogen specific protease cleavage site with a flexible Glycine-Serine motif. The targets for the antibodies and the cleavage sequences are unique for each of the three pathogens to impart high specificity.

**Fig.1d.**
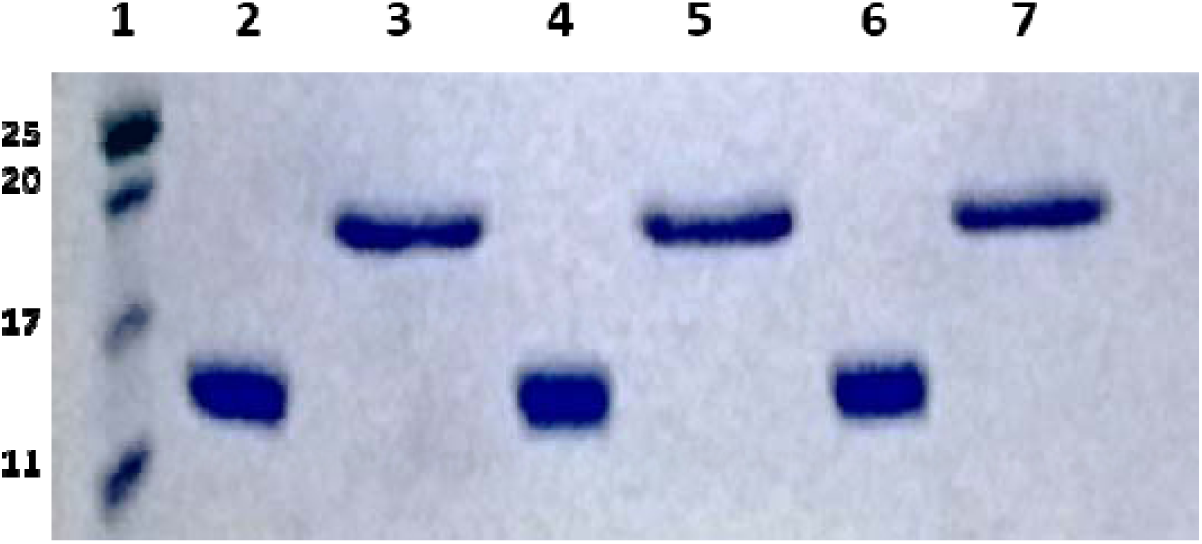
SDS Page analysis of purified VHH and the corresponding AbTids. They were expressed in *E. coli* with histidine tag and purified by affinity chromatography on a Nickel-NTA sepharose column to obtain molecules of high purity. lane 1 - marker, lane 2 and 3 – Candida CaC11h VHH and AbTid ABG 07, lanes 4 and 5 – Pseudomonas PsC23 VHH, and AbTid ABG 14, lanes 6 and 7 – Staphylococcus Sa C19 VHH and AbTid ABG 16.

#### 3.1.1 VHH surface targets

Two pathogen surface targets for VHH were chosen, enolase for *S. aureus* and *Candida sp.*, and C4-dicarboxylate transporter for *P. aeruginosa* after careful consideration of the abundance and specificity on the pathogen surface.

##### C4 dicarboxylate transporters

*P. aeruginosa* is present in microaerophilic conditions in the host, has a capacity to adapt to different physiological conditions and utilise various carbon sources for energy, particularly C4-carbon metabolites like malate, fumarate, and succinate. This is due to the glyoxylate shunt, a metabolic pathway bypassing the TCA cycle which is absent in the human host. Utilizing this pathway, C4-sugars are imported using the C4-dicarboxylate transporters. This metabolic pathway plays an important role in *P. aeruginosa* persistence and drug resistance at the site of infection and was chosen as a target for development of VHH (39–46).

##### Enolase

Enolase, a metabolic enzyme that is present in pathogenic organisms like *C. albicans, S. aureus* and *P. aeruginosa* was chosen for both *C. albicans* and *S. aureus.* Enolase of *C. albicans* has a dual role: as glycolytic enzyme and is responsible for colonization in small intestinal mucosa (35,36). Enolase is required for conversion of 2-phosphoglycerate to phosphoenolpyruvate and acts as a plasminogen receptor as well. Infection in *C. albicans* and *S. aureus* is initiated by adherence to the host cell where the enolase interacts with host plasminogen that enhances plasmin activity of the pathogens by promoting the degradation of extracellular matrix (37,38). Since enolase is also present in *Homo sapiens*, homology search was done to identify the unique domains of the pathogens that are not present in humans (an important prerequisite to develop antibodies with high specificity). Sequence alignment revealed the homology of enolase between Candida and Staphylococcus to be 52.90%, Staphylococcus and humans 49.65% whereas the homology between Candida and human was 63.59% (Table 1). The VHH fragments were carefully selected against the non-homologous regions to ensure pathogen specificity and to reduce the chances of off-target activity to the host human enolase.

#### 3.1.2 Cleavage sites

The shortlisted VHH hits were attached to the AMP histatin by a pathogen-specific cleavable linker. These linkers, specific for proteases that aid in initial attachment and subsequent tissue penetration and play an important role in establishment of acute and chronic infections (47). They are abundant in the tissue invading stages and are expected to be present universally in all strains of the three pathogens. The following proteases for each of the pathogens were chosen for designing cleavable linkers for this study.

##### P. aeruginosa

Pseudolysin, the extracellular endopeptidase of *P. aeruginosa* commonly called ‘Pseudomonas elastase’ or ‘Elastase B’ is a zinc-dependent metalloprotease that belongs to the M4 thermolysin-like family (48). It causes deterioration of the human arterial elastic laminae during systemic infections (49).

##### S. aureus

The serine protease known as ‘V8’ protease belongs to glutamyl endopeptidase family, is a virulence factor and has three forms: Staphylococcus serine protease (Ssp A, Ssp B and Ssp C). Ssp A is one of the immune suppression factors present in variety of innate immune system suppression pathways and can also degrade complement components, induce extracellular matrix structural damage, and promote vascular leakage (49,50). It promotes the engulfment of neutrophil by microphages or apoptosis of neutrophils and inhibit neutrophil chemotaxis (51).

##### C. albicans

Secretory aspartyl proteases (Sap) are required for tissue invasion and colonisation due to their broad substrate specificity. Target of Sap proteases include enzymes from macrophages, keratin, proteinase inhibitors, collagen, mucin, and enzymes of the blood-clotting cascade (52).

#### 3.1.3 Linker design and activation assay

The protease cleavage sites were designed from MEROPS database (https://www.ebi.ac.uk/merops/) and were placed between the AMP and the VHH by flexible Glycine-Serine linkers of two different sizes (G4S)3 and G4S to allow for structural flexibility and ensuring solubility of the complex. As the size of the AbTids (20 kDa) is larger than the VHHs (14 kDa) (Fig. 1d), activation was expected to be accompanied by lowering the size to 14 kDa due to release of the antimicrobial peptide by the pathogen proteases that was detected by SDS PAGE assay.

### 3.2 Cloning and expression of pathogen surface targets and specificity analysis

To generate VHH hits against the two enolases, we had to carefully select VHHs without any cross reactivity to each other as well as with the human enolase. To avoid risk of cross reactivity of the AbTid with enolases in different organisms, we expressed the 48 kDa *Candida sp* (Fig. 2d C) and *S. aureus* (Fig. 2d A) enolases in *E. coli* and used them as an immobilised antigen by bio-panning to isolate pathogen specific, non-cross-reacting VHH fragments from the CIL-Ca and CIL-Sa libraries. Human enolase (Recombinant human enolase-1, ENO1 Human, PROSPEC protein specialists, Israel) which was 47.1 kDa (434 amino acids) protein was used as a negative control (Fig 2d B and D) for the western blots and only those VHH clones that bound to the pathogen and not to the human enolases were selected as specific hits to *S. aureus* and *C. albicans* (Fig. 2d A and 2d C) and used for the development of AbTids.

**Fig 2.**
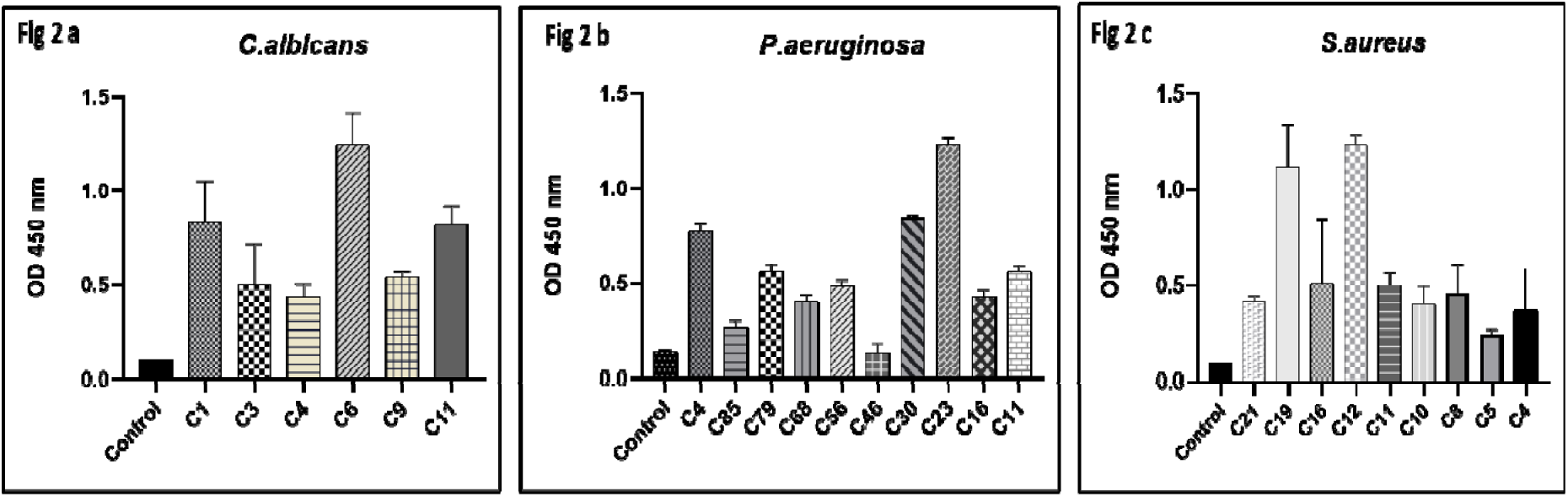
**a,b,c.** Hit selection using whole cell phage-ELISA with supernatant of randomly selected clones from the pathogen-specific libraries. The pathogens were fixed on the plates and incubated with the supernatant of *E. coli* clones containing the full-length VHH in pADL-23c vector induced with IPTG. The binding intensity was measured by ELISA using mouse anticamel HRP antibody. A more than 4-fold increase compared to control was identified as a strong binder and *C. albicans* C6 (Fig. 2a), *P. aeruginosa* C23 (Fig. 2b) and *S. aureus* C19 (Fig. 2c) were progressed further.

**Fig. 2d.**
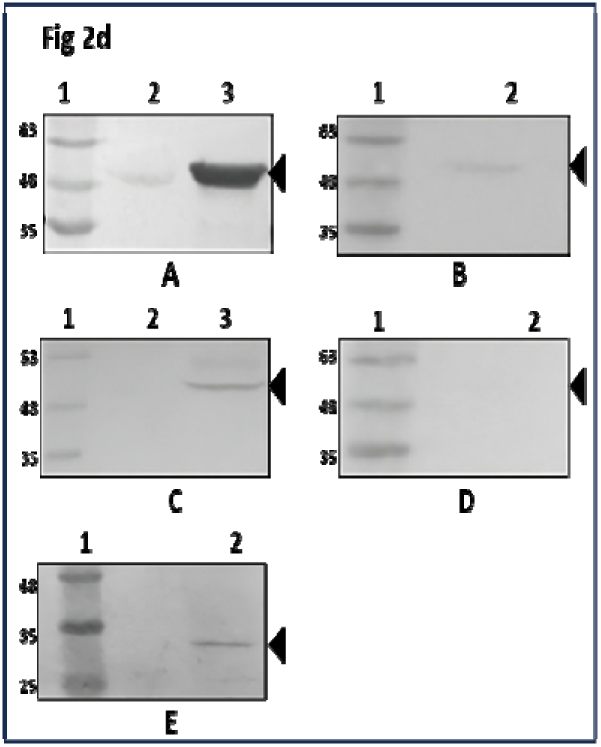
Western blot analysis of recombinantly produced pathogen surface targets probed with different AbTids. The targets C4 di-carboxylate transporter for *P. aeruginosa*, enolase for *C. albicans* and *S. aureus* were expressed in *E. coli* and probed with the purified AbTids ABG 07, ABG 14 and ABG 16. 1 - *S. aureus* AbTid ABG16. lane 1 – Marker, lane 2 – Human enolase, lane 3 – Staph. Enolase. 2 - *S. aureus* AbTid ABG16 lane 1 – Marker, lane 2 – Candida enolase. 3 - *C. albicans* AbTid ABG07 lane 1 – Marker, lane 2 – Human enolase, lane 3 – Candida enolase. 4 - *C. albicans* AbTid ABG07 lane 1 – Marker, lane 2 – Staph enolase. 5 - *P. aeruginosa* AbTid ABG14 lane 1 – Marker, lane 2 – C4 di-carboxylate transporter.

For *P. aeruginosa*, a 33 kDa subunit of the C4-dicarboxylate transporter was extracted out by PCR from the *P. aeruginosa* BAA 2108 (Carbapenem resistant) strain and recombinantly expressed in *E. coli* in pET 26a (Fig. 2d E) and used to isolate VHH against using the CIL-Pa library

### 3.3 Construction of VHH libraries and isolation of hits against surface targets of *P. aeruginosa, S. aureus and C. albicans*

All the three immunised libraries were constructed using the same procedure. For *P. aeruginosa,* a library (CIL-Pa) of 2×10^9^ clones, for *S. aureus*, a library (CIL-Sa) of 5×10^8^ clones and for *C. albicans*, library (CIL-Ca) of 1×10⁸ clones were generated with a diversity of > 90% for each.

Hits were isolated from the corresponding libraries against recombinantly produced surface targets and analysed by ELISA. 10-hits against C4-dicarboxylate transporter of *P. aeruginosa*, 9-hits against enolase of *S. aureus*, and 6-hits against enolase against *C. albicans* were shortlisted for further analysis by ELISA. The hits were sequenced and the ones containing the intact VHH with all the framework and CDR regions were progressed to the next stage of development by joining them to the AMP and the cleavable linkers. From CIL-Pa library, the clones C4, C23, C10 were analysed further, out of which C23 had an intact VHH sequence with discernible framework and CDR regions that also showed good expression. (Fig. 2b). For CIL-Sa, shortlisted clones were C19, C12, and C19, and were taken for further development as they contained a full-length gene that did not cross react with the human enolase (Fig. 2c). For CIL-Ca, the shortlisted clones were C1, C6, C11, out of which C11 was chosen for further development based on the above criteria (Fig. 2a). Two residues (Glu 49 to Gly and Arg 50 to Leu) were changed to humanise this molecule and the clone was designated as ‘C11h’. The AMP and the linker sequences were joined to the Pa C23, Sa C19 and Ca C11h to generate the three different AbTids used in this study – ABG 07 for *C. albicans*, ABG 14 for *P. aeruginosa* and ABG 16 for *S. aureus*.

The AbTids, they were then analysed for specificity to the designated targets by ELISA using the recombinantly produced antigens. Extreme specificity with no cross reactivity was seen in case of all the three AbTids indicating successful development of the molecules with specificity to its designated targets (Fig. 2e).

**Fig. 2e.**
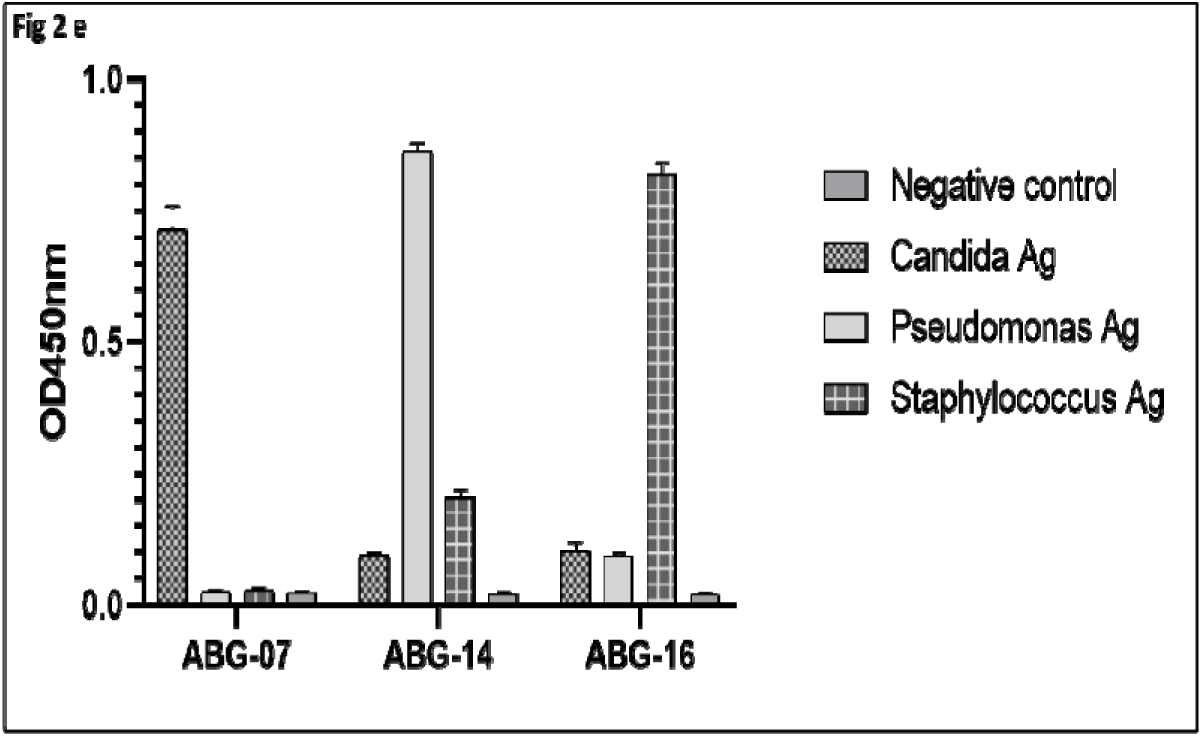
ELISA for binding specificity and cross reactivity of the AbTids against the purified target antigens. The purified AbTids (ABG 07, ABG 14 and ABG 16) were fixed on the plates and incubated with histidine tagged antigens C4 di-carboxylate transporter for *P. aeruginosa* and enolase for both *C. albicans* and *S. aureus* followed by detection with secondary antibody (anti HIS-HRP).

### 3.4 Activation of AbTids by pathogen-protease mediated release of AMP

The protease cleavage sites were chosen based on the possibility of their presence in all the strains of each of the pathogens. The functionality of the cleavage sites - Las B for *P. aeruginosa*, SspA for *S. aureus* and Sap for *Candida sp.*, were used for activation (Fig. 1c). The AbTids were incubated with the supernatant of the pathogens grown in Mueller-Hinton medium with added FBS to mimic the physiological conditions as closely as possible. In all the three AbTids, cleavage was initiated after 2-hours of incubation that continued for 4-hours, and complete cleavage was seen, after 6-hours with ∼14 kDa VHH band released from the AbTid (Fig. 1d). The activation of the AbTid by cleavage was very specific for the pathogens in both the strains tested and there was no cross cleavage by different pathogens. When *Pseudomonas* AbTid ABG 14 was incubated with *C. albicans* and *S. aureus* supernatants, cleavage was not seen (Fig. 3b). Likewise, cleavage was not observed in *Candida* AbTid ABG 07 when incubated with *P. aeruginosa* and *S. aureus* (Fig. 3a) and same result was seen when *Staphylococcus* AbTid ABG 16 when incubated with *P. aeruginosa* and both species of *Candida* (Fig. 3c). Proper selection of the cleavage sequences resulted in extreme specificity that ensured minimal off-target activity, a prerequisite for non-traditional antibiotic development.

**Fig. 3:**
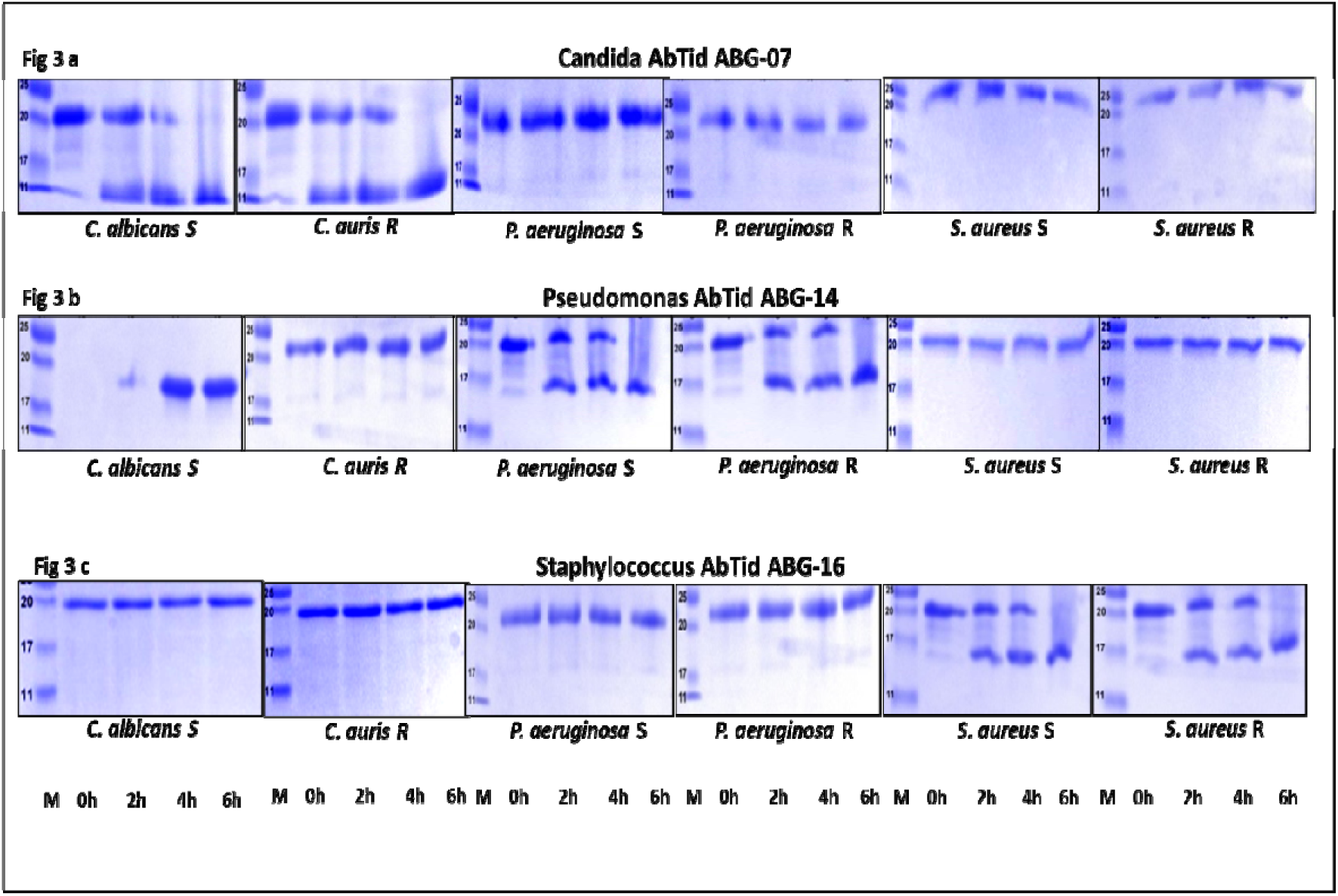
SDS PAGE analysis for the specificity of cleavage of AbTids in supernatants containing pathogen proteases. The three pathogens were grown to OD_600_1 and the AbTids (1 mg) incubated with the supernatants for 2, 4 and 6 hr. Two strains were used in each case, one drug sensitive (S) and the other drug resistant (R). *C. albicans* is the S strain and *C. auris* that is naturally multidrug resistant is R. Samples were analysed for their activation by the release of the peptide indicated by their cleavage pattern. From L to R Lane 1-Marker, Lane 2: 0 hr., Lane 3: 2 hr., Lane 4: 4 hr. and Lane 5: 6 hr.

**Fig. 3a.** Cleavage assay of anti-Candida AbTid ABG 07 with *C. albicans*, *C. auris*, *P. aeruginosa* S (ATCC 27853) and *P. aeruginosa* R (MCC 50483), *S. aureus* S (MCC 511005), (vi) S aureus R (MCC 50811). Fig 3b. Cleavage assay of anti-Pseudomonas AbTid ABG 14 with the pathogens in the same order and Fig 3c. Cleavage assay of anti-Staphylococcus AbTid ABG 16 with the pathogens in the same order. Specificity of the cleavage is demonstrated by the presence of two bands (partial digestion of the AbTids) in 2 hr. and 4 hr. and finally a single band in 6 hr. when complete cleavage and activation occurs.

**Fig. 3d.**
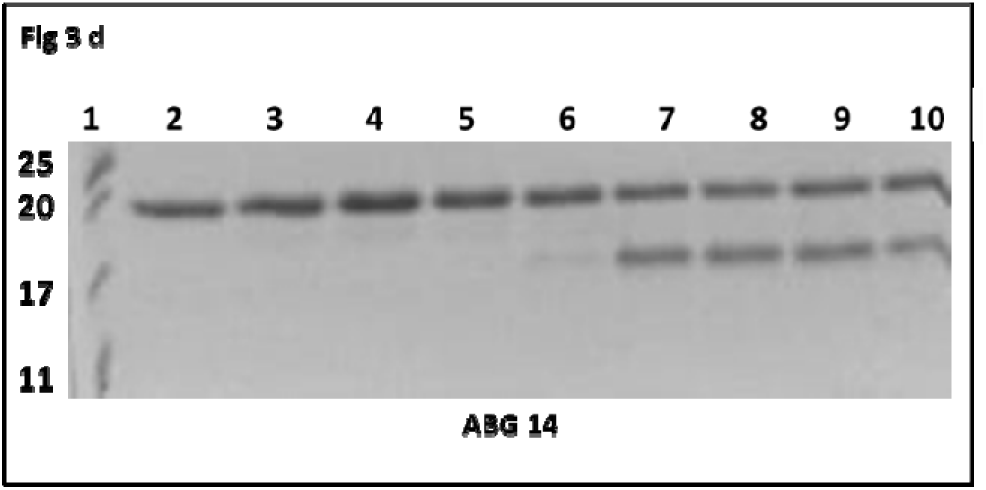
Relationship of cleavage activity of AbTids and pathogen count. SDS PAGE analysis for cleavage of ABG 14 in 2 hr. with increasing CFU mL^-1^ of carbapenem resistant *P. aeruginosa* MCC 50483. *P. aeruginosa* was grown to OD600and dilution adjusted to different cell counts. The supernatant was harvested by centrifugation and incubated with ABG 14. Lane 1 – Marker, lane 2 – ABG 14, lane 3 –10^2^ CFU, lane 4 –10^3^ CFU mL^-1^, lane 5 –10^4^ CFU mL^-1^, lane 6 –10^5^ CFU mL^-1^, lane 7 –10 6CFU mL^-1^, lane 8 –10^7^ CFU mL^-1^, lane 9 –10^8^ CFU mL^-1^, lane 10 –10^9^ CFU mL^-1^.

**Fig. 3e.**
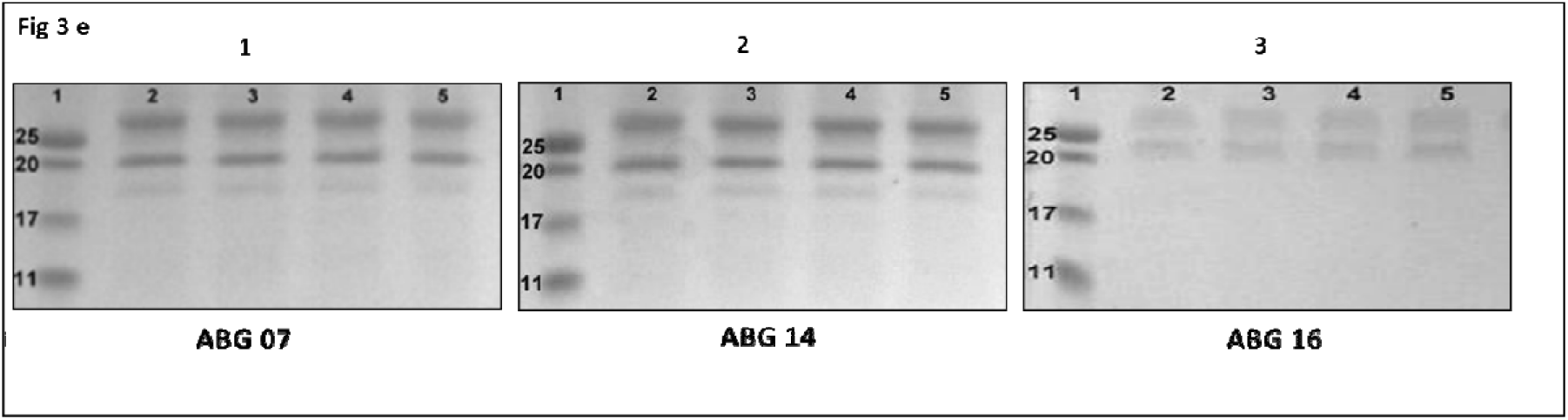
SDS PAGE analysis of stability of the three AbTids in human plasma. Plasma stability assay was performed using complement inactivated human plasma that was incubated at 37°C for 0, 2, 4 and 24 hr. with 4.76 μM of three different AbTids. 1. Anti-candida AbTid ABG 07, 2. Anti-Pseudomonas AbTid ABG 14 and 3. Anti-Staphylococcus AbTid ABG 16.

We also checked the dose of the protease required for cleavage mediated activation by incubating the Pseudomonas AbTid ABG 14 with increasing concentrations of supernatant by increasing the number of *P. aeruginosa* MCC 50428 (Fig. 3d) and analysing the result after 2-hours, during which activation of the molecule by protease cleavage was initiated. Cleavage was seen from 10⁵ CFU mL^-1^ and was consistently maintained till 10⁹ CFU mL^-1^ . At 10⁶ CFU mL^-1^ bacteria, the cleavage was completed in 2 hr. and the molecule exerted its full bactericidal potential. 10⁵ CFU mL^-1^ concentration is commonly found in patients with sepsis as well as abscesses (54), and this class of molecules can be expected to be exert a bactericidal effect in physiological conditions.

### 3.5 AbTids neutralise the pathogens specifically with minimal cross reactivity

The MIC of ABG 07, ABG 14 and ABG 16 on a sensitive and a resistant strain of all the three pathogens were evaluated *in vitro*. The pathogens were incubated with varying concentrations of both the VHH and the AbTids and their microbicidal effect was evaluated by monitoring their growth for 16 hours, quantification of the CFU by dilution plating on the Muller-Hinton agar plates.

#### 3.5.1 Efficacy

A good dose response neutralising effect was seen in the case of the VHHs and the AbTids, indicating specific activity. 90% to 95% killing was observed with Candida CaC11 VHH at concentration of 75 and 100 µg mL^-1^ and similar killing was observed with the corresponding ABG-07 AbTid at concentration of 5 and 10 µg mL^-1^ against both *C. albicans* and *C. auris*. The Candida AbTid were 10-15 times more efficacious than the VHH fragment alone in terms of bactericidal activity (Fig. 4a). *Candida auris* is resistant to all major classes of antifungals and both the VHH and ABG 07 neutralises both *C. albicans* and *C. auris* effectively, indicating their efficacy against the drug-resistant Candida. As the target and the cleavage enzymes are present in other Candida, this neutralising activity can be expected against other Candida species as well, making ABG 07 a new anti-Candida hit.

**Fig. 4:**
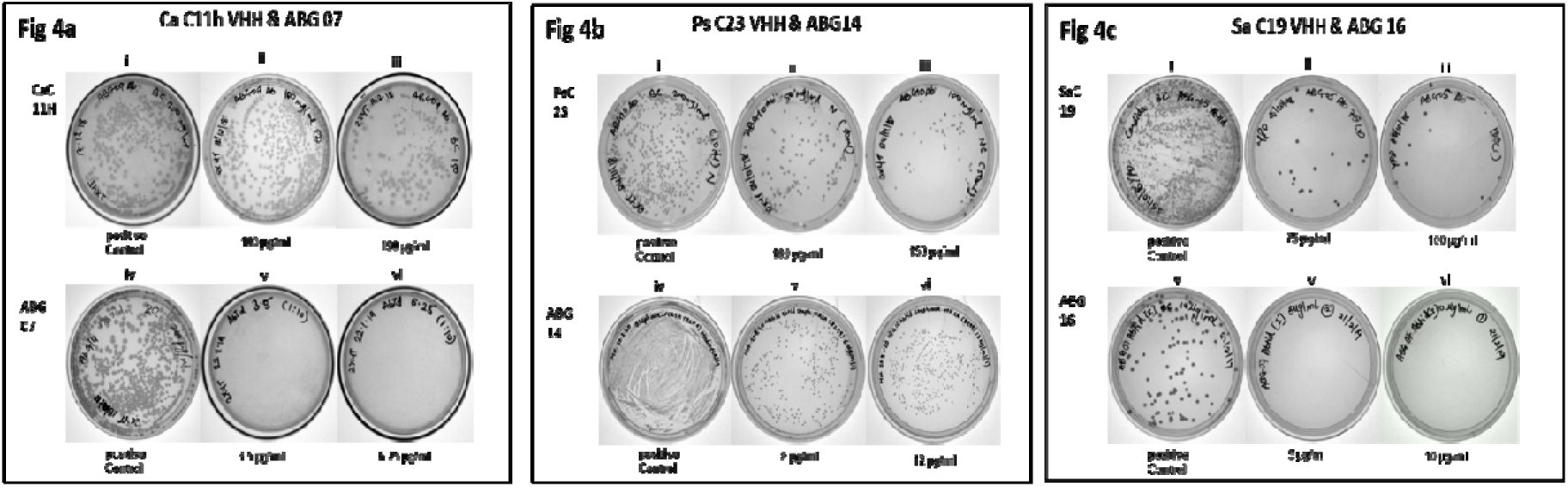
**a,b,c.** Determination of the efficacy of the VHHs and AbTids by microbiological spread plating. The pathogens were grown to OD_600_1 diluted to 50,000 CFU/mL and incubated with the presence of the VHH (75-150 μg mL^-1^) and AbTids (5-25 μg mL^-1^) overnight, plated in MH agar and the CFU mL^-1^ quantitated. Positive controls are plating without any drug treatment. a. Effect of C11h VHH and ABG 07 AbTid on *C. albicans*. b. Effect of PsC23 VHH and ABG 14 on multidrug resistant *P. aeruginosa* BAA 2108. c. Effect of SaC19 and ABG 16 AbTid on Methicillin resistant *S. aureus.* MCC 50811.

**Fig. 4:**
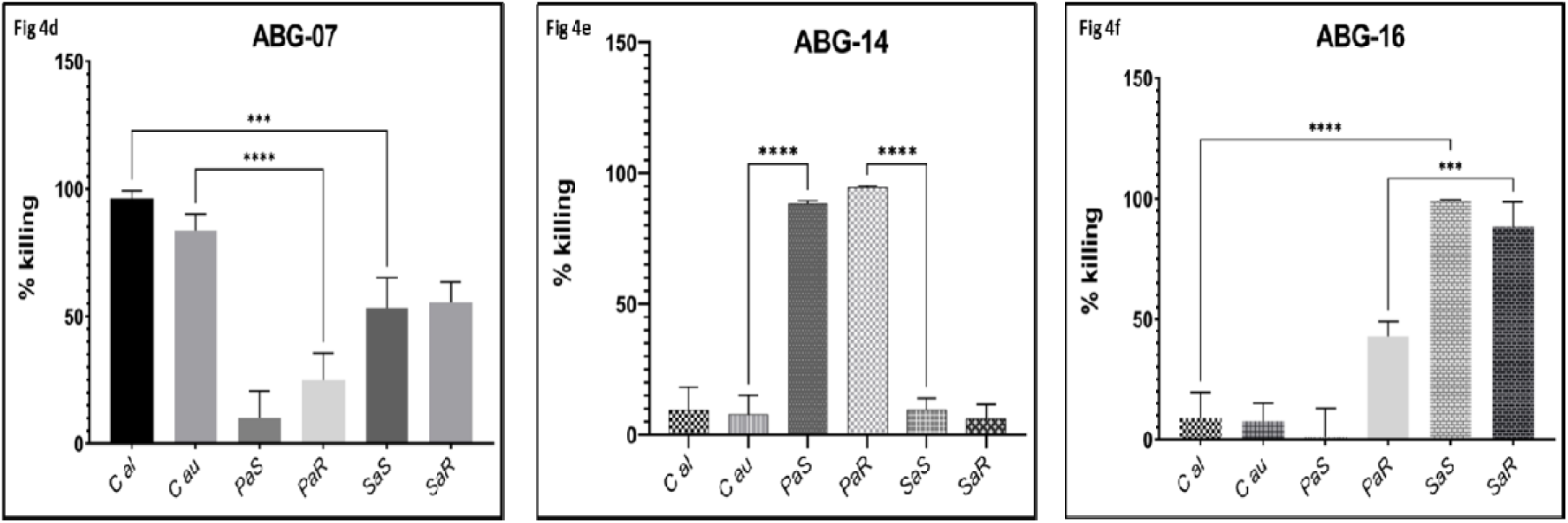
**d,e,f.** Specificity of the AbTids as determined by percentage killing on two different strains of the three pathogens. The pathogens were diluted to 50,000 CFU, AbTids were administered at a dose of 25µg mL^-1^ incubated overnight and plated. The sensitive pathogens are *C. albicans* (Cal), *P. aeruginosa* ATCC 27853 (PaS) and *S. aureus* MCC 511005 (SaS) and the resistant pathogens are *C. auris* (Cau), *P. aeruginosa* BAA 2108 (PaR) and *S. aureus* MCC 50811 (SaR). Statistical analysis was performed consisting of one-way ANOVA and Tukey post hoc test (set threshold values: p < 0.05 and confidence level = 95%. a. Specificity of the anti-Candida AbTid ABG 07 on sensitive and resistant strains of *Candida sp., Pseudomonas sp.* and *Staphylococcus sp*. b. Specificity of the anti-Pseudomonas AbTid ABG 14 on sensitive and resistant strains of *Candida sp., Pseudomonas sp.* and *Staphylococcus sp*. c. Specificity of the anti-Staphylococcus AbTid ABG 16 on sensitive and resistant strains of *Candida sp., Pseudomonas sp.* and *Staphylococcus sp*.

85% to 90% killing was observed with anti-Pseudomonas PsC23 VHH at concentrations of 100 and 150 µg mL^-1^ and at 3.5 and 6.25 µg mL^-1^ for the corresponding AbTid ABG 14 against multidrug resistant *P. aeruginosa* ATCC BAA 2108 and the reference sensitive *P. aeruginosa (*ATCC 27853), indicating that the AbTid was 25-30 times more effective than the VHH alone (Fig. 4b) and was identified as a hit against drug resistant forms of *P. aeruginosa*.

About 80% to 90% killing were observed in anti-Staphylococcus SaC19 VHH with concentration of 50 and 100 µg mL^-1^ and similar levels of killing was observed with corresponding ABG 16 AbTid with concentration of 6 and 12 µg mL^-1^ against methicillin resistant *Staphylococcus aureus* MCC 50811 (MRSA) and methicillin sensitive MCC 51005.The AbTid was seen to be 90% more effective in neutralising the pathogen as the MIC 90 dose dropped from 100 µg mL^-1^ to 12 µg mL^-1^ in both the strains tested (Fig. 4c). ABG 16 is effective against MRSA and can be considered as a hit suitable for further development.

Both the new sets of antibacterial hits, the VHH and their corresponding AbTids were found to be bactericidal irrespective of the drug resistance profile of the pathogens. A similar effect was seen in *P. aeruginosa* BAA 2108 (multi-drug-resistant strain) and methicillin resistant *S. aureus* MCC 50811 and multidrug resistant *C. auris* (data not shown). These results clearly demonstrate that the microbicidal action of the antibody-drug conjugate is more efficacious than the corresponding VHH alone irrespective of the resistance profile of the pathogens.

#### 3.5.2 Specificity

The specificity of AbTids was due to the VHH precisely targeted to the pathogen surface target as well as the protease dependant cleavage sequences a few amino acids long; that ensures an effective attachment followed by cleavage mediated release of the AMP resulting in activation of the molecule. We checked the specificity of the three AbTids by checking their efficacy against two strains of the target pathogens as well as by cross reactivity studies to the other non-target pathogens.

The anti-Candida ABG 07 AbTid demonstrated 80-90% efficacy not only in both *C. albicans* MTCC 227 and *C. auris* ATCC MYA 5002, but some consistent cross reactivity (40%-50%) is seen with both the *S. aureus* strains MCC 50811 (Methicillin resistant) and MCC 51005 (Methicillin resistant), that is not statistically significant (Fig. 4d). This effect is possibly due to the common target enolase for both the pathogens, even though the VHHs were not found to be cross reactive when checked against purified antigens (Fig. 2e). There is negligible cross reactivity with the *P. aeruginosa* strains (about 15-20%), but the overall specificity of ABG 07 is significant for Candida.

The specific action of anti-Pseudomonas ABG 14 AbTid against both the carbapenem sensitive and resistant strains of *P. aeruginosa* was the most significant, with the percentage killing by ABG 14 being around 80-90%. No cross reactivity was observed with the two other pathogens possibly because there was no homology between the targets (Fig. 4e).

For the anti-Staphylococcus molecule ABG 16, > 90% killing was seen in both the methicillin sensitive and resistant strains of Staphylococcus, making it specific for *S. aureus*. Slight cross reactivity (40%) was also observed with one of the strains *P. aeruginosa* BAA 2108, which is multidrug resistant. For rest of the two pathogens in this study, there was negligible cross reactivity in the range of 5-10% (Fig. 4f).

There was a statistically significant difference between cross-reactivities as determined by one-way ANOVA [(ABG 07 F (5,12) = 40.40, p < 0.0001), (ABG 14 F (5,12) = 186.7, p < 0.0001), (ABG 16 F (5,12) = 72.35, p < 0.0001)]. A Tukey’s HSD post hoc test revealed that the specificity of AbTids against the target pathogens was statistically significant (α < 0.0001) while reactivity with other pathogens showed no statistical significance at threshold values of α < 0.05 and confidence level of 95%.

#### 3.5.3 ABG 14 specifically neutralises carbapenem resistant strain *P. aeruginosa*, destroys biofilms and has no effect on beneficial gut bacteria

The effect of ABG14, PsC23 and meropenem were evaluated against the reference strain ATCC 27853, and the meropenem resistant strain MCC 50428 of *P.aeruginosa*. We performed time-kill assay with 6.25 µg mL^-1^ of ABG 14, 25 µg mL^-1^ of PsC23 and 8 µg mL^-1^ of meropenem and monitored the number of viable cells over the period 2, 4, 8, and 24 h (Fig. 5e). and the most pronounced effect was observed within 2 h after which the bacteria recovered and a gently upward sloping curve was observed for both PsC23 and ABG 14. Although this result was not reflected in the dilution plating results, due to a six log difference between the control, it is possible that the amount of the molecules somehow became limiting or the action was bacteriostatic initially that was overcome by the rapidly multiplying bacteria. To confirm this possibility, we administered PsC23 and ABG 14 every two hours and a complete bactericidal effect was observed and no recovery of the pathogen population was seen till 24 hrs (results not shown). The neutralisation activity and cross-activity of purified ABG 14 against probiotic bacteria was then evaluated by monitoring the growth of the two strains of *P. aeruginosa* (ATCC 27853 and multidrug resistant BAA 2108) and six other bacteria (four *Lactobacillis sp.* and *E. coli* and *B. subtilis*) after incubating them with the AbTid (12.5 µg mL^-1^) overnight. ABG 14 demonstrated a neutralizing activity against both the strains of *P. aeruginosa* but did not significantly affect the growth of the other tested bacteria (Fig. 5a). This indicated that the ABG 14 was very specific in terms of binding and neutralization abilities due to its dual mode of action. We next determined the efficacy of ABG 14 against *P.aeruginosa* biofilms, that would normally be present in physiological conditions. The small size ensured good substrate penetration and the anti-biofilm properties of the AMP histatin (55) the ABG 14 (6.25 µg mL^-1^) resulted in interference with the biofilm formation in the multidrug strain BAA 2108 as indicated by the lack of colour reaction to 0.1% crystal violet (Fig. 5b).

**Fig. 5:** Characterisation of *P. aeruginosa* AbTid ABG 14.

**Fig. 5a:**
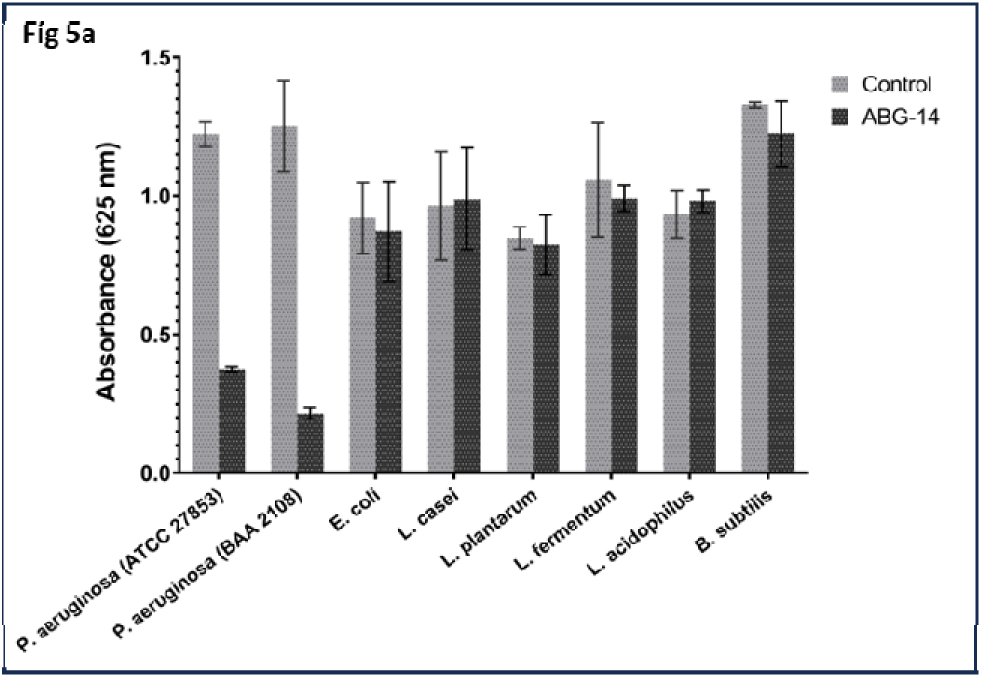
Microbicidal action of ABG 14 against *P. aeruginosa* ATCC 27853 and multidrug resistant BAA 2108 and commensal bacteria of the Lactobacillus group, *E. coli* and *B. subtilis*.

**Fig. 5b:**
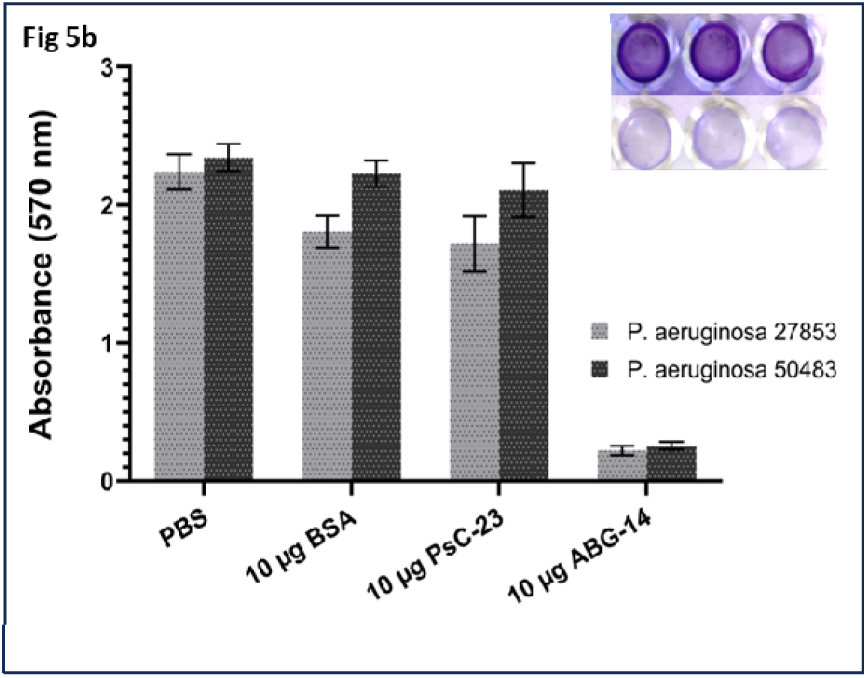
Effect of ABG 14 on biofilm formation in *P. aeruginosa* 27853 and MCC 50428 by quantitation of crystal violet stain by dissolving in 95% ethanol. Inset: Visualisation (top no treatment, bottom: with ABG 14) of the crystal violet stained wells.

#### 3.5.4 AbTids are plasma stable and non-toxic to human cells

As the AbTids have been developed for parenteral use to target well entrenched pathogens in different parts of the body, a plasma stability assay was performed. To check the stability of the molecule in plasma where similar proteases might be present, all the three AbTids were incubated in human plasma for 24-hours and checked for their integrity on SDS-PAGE. No cleavage of any of the AbTids was observed till 24-hours (Fig. 3e 1,2,3), indicating that plasma proteases had no effect on the cleavage sites and subsequent stability of the molecules; the molecules were activated only in the presence of the pathogen proteases, an important druggable property. The toxicity of ABG 14 was tested on human cell line HEK 293 by MTT assay (Fig 5c) and no significant toxicity was seen at 20 times the effective dose (125 µg mL^-1^).

#### 3.5.5 *P. aeruginosa* 27853 is unable to acquire resistance to ABG 14 but easily becomes resistant to meropenem

When *P. aeruginosa* 27853 that is susceptible to meropenem at a concentration of 1 µg mL^-1^ was grown in a sublethal dose of 0.5 µg mL^-1^, a negligible increase in growth was seen in the first three generations. However, by the end of the next three generations, a threefold increase in growth was observed followed by an equivalent increase in the next three generations (Fig. 5d), indicating that the pathogen slowly became resistant to the drug. This dose had no effect on the carbapenem resistant MCC 50428 strain and the bacteria showed a robust growth in all the three generations. When a sublethal dose of 3.125 µg mL^-1^ of ABG 14 was administered to both the strains of *P. aeruginosa*, there was no appreciable growth across the nine generations indicating that the bacteria were incapable of acquiring resistance to the AbTid. Colony counting also showed a gradual increase in the number of *P. aeruginosa* 27853 in meropenem but no such increase was seen in ABG 14 (data not shown). Results of this experiment indicates that *P. aeruginosa* easily develops resistance against small molecule antibiotic like meropenem but is unable to do the same against a complex biological like ABG 14. AbTids are thus expected to be resilient to mutation-based resistance.

#### 3.5.6 ABG 14 clears systemic infection of carbapenem resistant *P. aeruginosa* from neutropenic mouse

Before advancing the ABG 14 to *in vivo* studies, its druggable properties were evaluated. It was found to be stable in human serum for 24 hr. and did not have a toxic effect on HEK 293T cells even at 100 µg mL^-1^ as evaluated by MTT assay (Fig. 5c) To evaluate the *in vivo* efficacy of ABG 14, its effect was studied in BALB/c mice with systemic infection of carbapenem resistant strain of *P. aeruginosa* MCC 50428. Various treatments with ABG 14 and meropenem were administered intravenously BID as mentioned in Table 2

. Heparinised blood was withdrawn from the tail vein after 24 hr. and the pathogen load was counted by plating. High bacterial counts were seen in mice administered with 5 mg kg^-1^ body weight dose of meropenem. Treatment of ABG 14 alone reduced the colony counts by 8-logs to 10-logs compared to control mice without any treatment (Fig. 5f, inset). Uninfected mice had no detectable pathogens and mice treated with ABG 14 showed no mortality till 72 hr, but infected mice treated with meropenem perished with 3-4 days with a similar effect seen with mice without treatment (Fig. 5f). The 5^th^ group of mice that were administered ABG 14 equivalent to 5 mg kg^-1^ body weight without infection showed a 75% survival for 72 hr. indicating non-toxic property of the VHH (data not shown).

**Fig. 5c:**
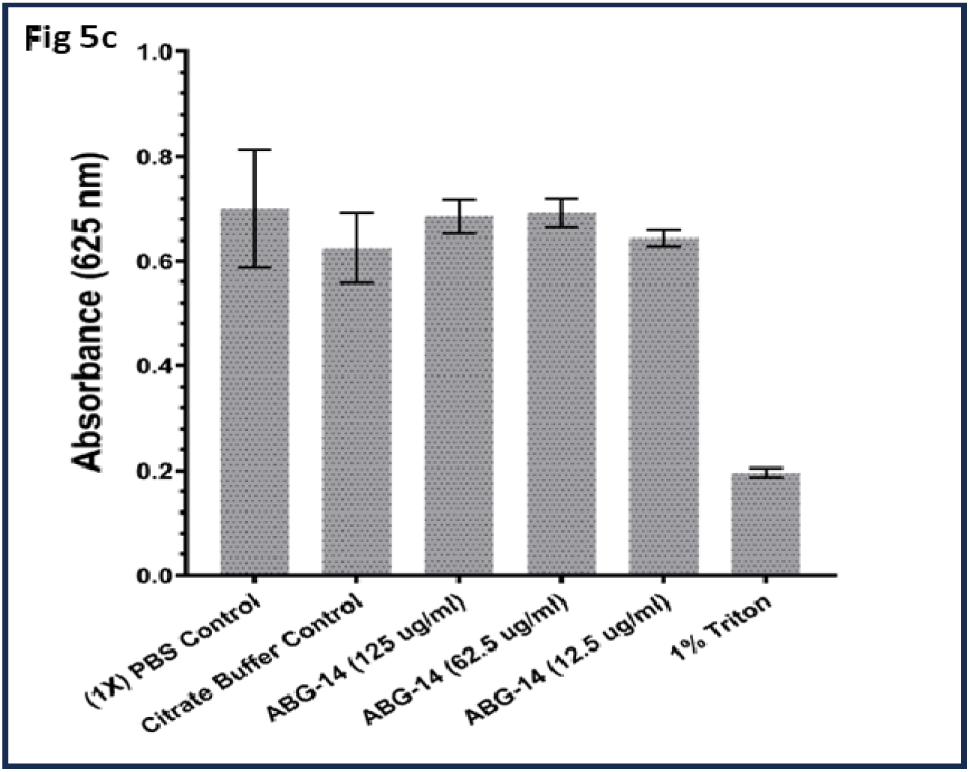
Cytotoxicity studies of ABG 14 on *P. aeruginosa* 27853. MTT assay was performed after incubating the human HEK 293K in different concentration of the AbTid ABG 14 for 24 hr. 1% Triton X-100 was used as a positive control.

**Table 2:** Experimental design to study the effect of ABG14 in carbapenem resistant P. aeruginosa MCC 50428 infected BALB/c mice.

| Group | Treatment | Dose (mg kg<sup>-1</sup>) |
| --- | --- | --- |
| 1 | No Infection | PBS, pH 7.2 |
| 2 | Infection + No treatment | PBS, pH 7.2 |
| 3. | Infection + ABG14 | 5 mg kg<sup>-1</sup> |
| 4 | Infection + Meropenem | 5mg kg<sup>-1</sup> |
| 5 | No infection ABG14 | 5 mg kg<sup>-1</sup> |

**Fig. 5d:**
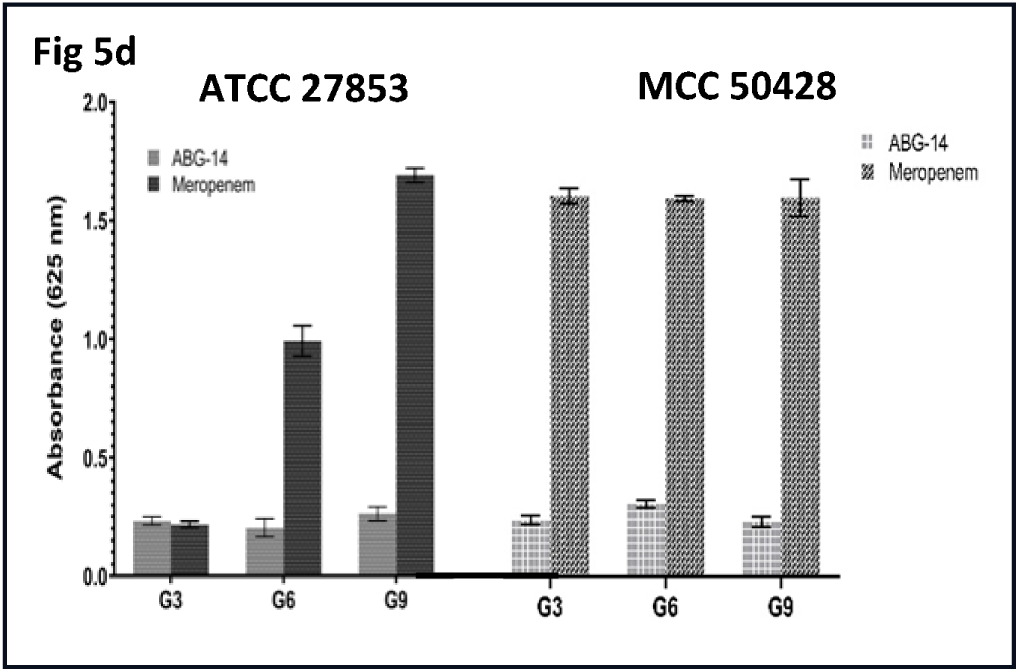
Adaptive resistance of *P. aeruginosa* 27853 (left) and carbapenem resistant MCC 50428 (right) to AbTid ABG 14 and meropenem. The pathogens were grown in sublethal doses of the two antimicrobials through nine generations (G1-G9) and the OD_600_ plotted at G3, G6 and G9.

**Fig 5e:**
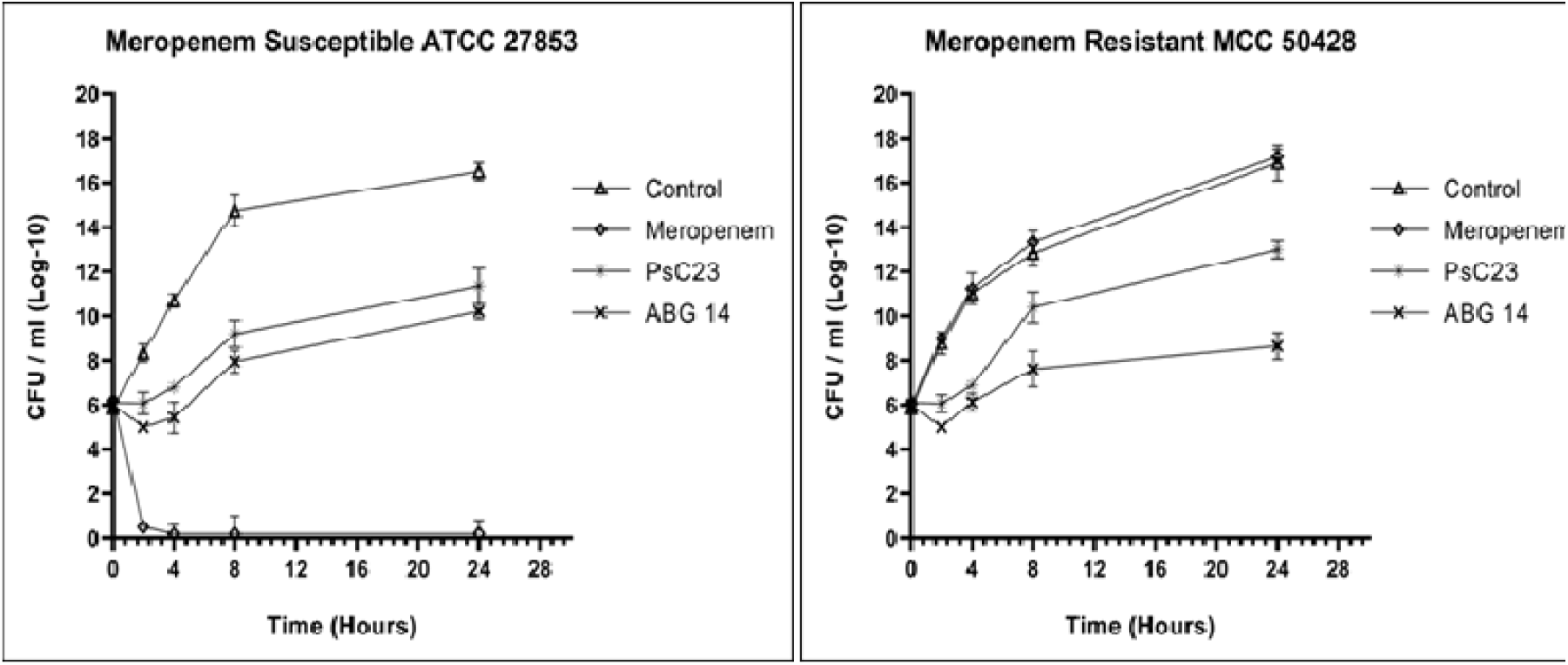
Growth kinetics of meropenem susceptible ATCC 27853 (left) and resistant MCC 50428 (right) *P. aeruginosa* in the presence of meropenem, PsC23 and ABG 14 over a period of 24 h.

**Fig. 5f:**
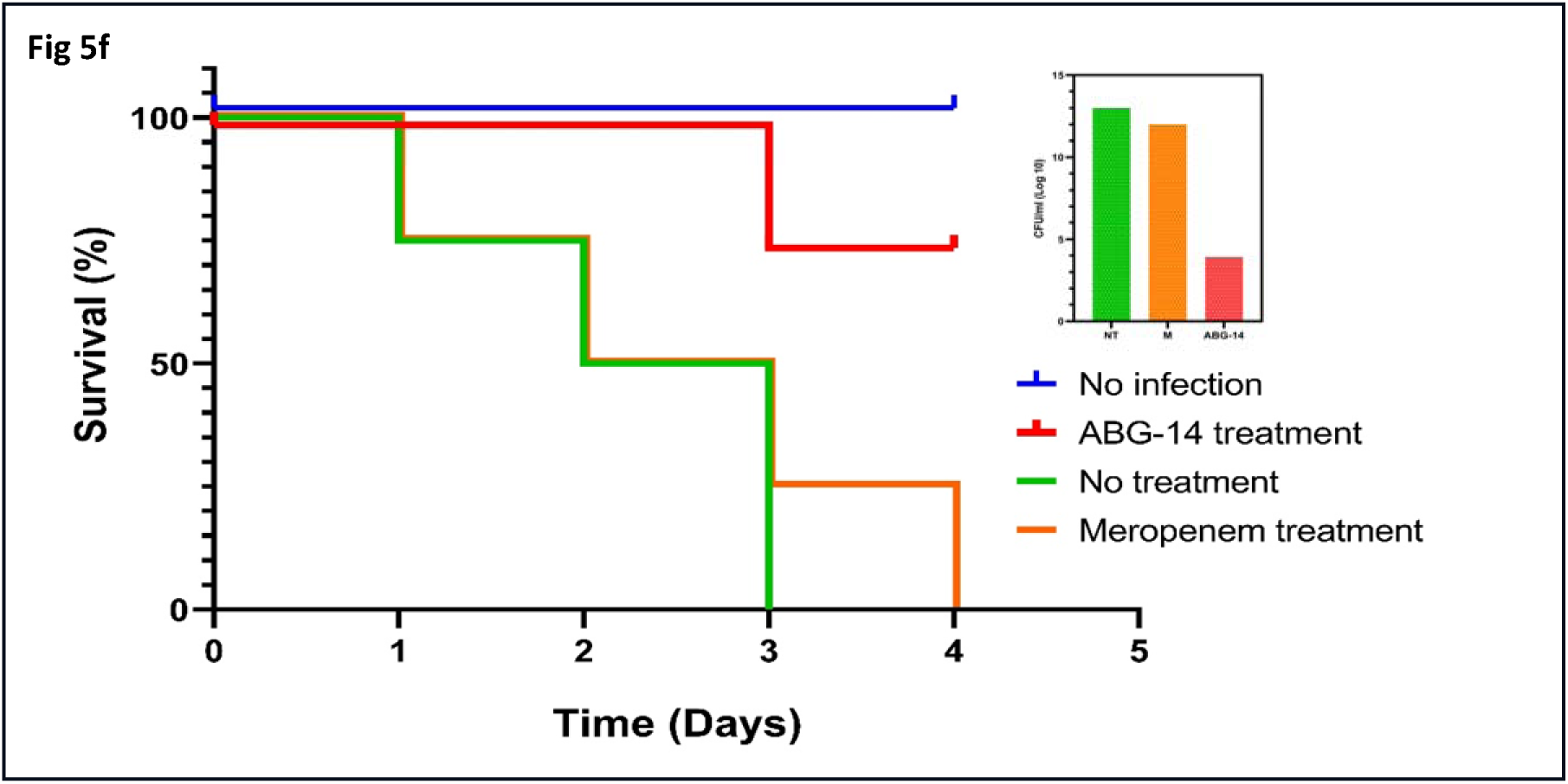
Survival curves neutropenic BALB/c mice infected with meropenem resistant *P. aeruginosa* MCC 50428 when treated with meropenem 5 mg Kg^-1^ (Group 4, orange), ABG 14 5 mg Kg^-1^ (Group 3, red), and without treatment (Group 2, green) and no infection (Group1, blue), monitored over a period of 72 hr. All mice administered ABG 14 with no infection (Group 5) survived the duration of experiment (data was not plotted). Fig. 5e **inset**: Bacterial load estimation in blood of BALB/c mice (Log 10 CFUmL^-1^) after 24 hr. of infection with carbapenem resistant *P. aeruginosa* MCC 50428 with and without treatment, with meropenem and ABG 14.

## 4. DISCUSSION

Multiple antibodies, mainly of human origin with opsonophagocytic and toxin neutralization mechanisms of action, have progressed to the clinical development stage for bacterial pathogens with a sizeable number targeting *P. aeruginosa* and *S. aureus* (56). The three approved antibody-based therapies in the market are based on toxin neutralization of *B. anthracis* and *C. difficile* (57). Antibody-drug conjugates against bacterial pathogens have been developed by Genentech and their human antibody-based *S. aureus* molecule Thiomab underwent Phase-I clinical trials (58), where intracellular localization of the molecule into the phago-lysosomal compartment and activation by cleavage in acidic pH were demonstrated. However, reports of progress could not be found from 2019 onwards, indicating that perhaps it might be undergoing modification(s) or further development had been terminated.

AbTids have been developed using a unique approach to negate the pitfalls of the antibody-drug conjugate methods tried earlier. As the pathogens in this study (except for *S. aureus*) were not known to produce toxins, the antibody fragments were chosen against surface targets. VHHs were used instead of full-length antibodies for the ease of tailoring and favorable druggable properties like low immunogenicity, stability, and short half-life. Due to their small size of 15 kDa, they can bind to cryptic antigens (19) and it is relatively easy to isolate a good number of hits against the target pathogens and winnow them by ELISA to enable the selection of non-cross-reacting antibodies in a short period of time.

To ensure coverage of different pathogen types based on their surface properties, we chose three different pathogens in the CDC and WHO priority list, gram-negative *P. aeruginosa*, gram-positive *S. aureus* and yeasts *C. albicans/auris*. Targets present on the surface of the pathogens having a critical role in their survival and pathogenicity were chosen to ensure the greatest possible coverage of the AbTids against all strains of the pathogens. For *P. aeruginosa,* a surface transporter discovered in our lab was used as a target. Although C4-dicarboxylate transporter is present in all the three pathogens and not in the human host, we used this target only for *P. aeruginosa* as it is a facultative anaerobe under physiological conditions where the transporter plays a crucial role in survival and drug resistance. The PsC23 antibody developed against this target was found to reverse resistance to carbapenems and β-lactams when used in combination (53), and the corresponding AbTid ABG 14 increased its efficacy by twenty-fold.

To highlight the significance of the pathogen specific cleavage for activation AbTids, we used the same target, enolase for *Candida sp*. and *Staphylococcus sp*. so that even if the antibody were to bind to the same target in both, activation could be linked to pathogen specific cleavage. We also made sure that both the VHH hits did not bind to and affect the activity of human enolase, with which they share 49.65% - 63.59% homology, an important prerequisite for them to be developed as a drug molecule with strong specificity. Fortunately, we were able to isolate extremely specific hits using our robust phage display antibody screening techniques as we were able to target more epitopes than a regular antibody due to its small size. Both the AbTids showed extreme binding specificity and did not cross-react with the purified antigens, although some cross neutralization (∼ 40 %) was seen in microbiology studies. It is possible that the Candida AbTid ABG 07 got loosely bound to some component of Staphylococcus enolase *in situ*, an effect that needs to be investigated further.

AbTids are cleavable antibody drug conjugate, and the linkers were designed for efficient payload release at the site of intended action. Numerous linkers like hydrazone linker, cathepsin B linker, glutathione sensitive and pyrophosphate linkers have been conjugated to biologicals by chemical conjugation (59). Payload (AMP) release was ensured by pathogen protease virulence factors that are extremely important for the pathogens and are present in medically important strains. As a loss of function via mutation would exert a tremendous genetic load resulting in a loss of pathogenicity, the chance of resistance loss due to mutation in a virulent strain would be low *ab initio.* Three unique protease cleavage sites were chosen that are mutually exclusive between the pathogens so that activation by specific cleavage would occur and their effect on the bacterial viability as well as cross reactivity (if any) could be studied. Proteases are present on the surface of the pathogens (48,60,61) and are regulated by operons. To keep them operational *in vitro*, we conducted the experiments in a medium supplemented in serum to mimic the physiological conditions. Adequate dose of the protease was achieved due to the close proximity of the AbTid to the pathogens due to VHH-antigen interaction. We performed a cleavage assay to check for the activation of the AbTids with the culture supernatant containing the shed proteases when pathogen numbers were in the range of 10^5^ CFU mL^-1^ that is generally found in the sputum or in organs like urinary bladder of infected patients (54). AbTids were found to be plasma stable as well and were activated only in the presence of the specific pathogen proteases. This extreme plasma stability and specificity at a dose of 10^4^ - 10^5^ CFU mL^-1^ combined with their activation in 2-4 hours attested to their druggable nature.

The cleavage sequences were linked to hydrophilic, flexible serine glycine linkers of different lengths to determine if that had any role in yield, solubility as well as in the efficacy of the molecules. All the variations turned out to be bactericidal, indicating that the linker length did not influence the antibacterial activity. The increase in the length of the linker from G3S to (G3S)4 did not interfere with the CDR3 region or the attachment of the VHH to the target and bactericidal action was not affected. The hydrophobic nature of all the molecules ensured that they could be produced as inclusion bodies in *E. coli*. Biologicals have been produced as inclusion bodies to increase the yield and to make the molecule commercially viable; yields in g L^-1^ are required which was achieved in the *Picchia* system (2-3 g L^-1^) in the soluble secretory form (data not shown).

Conjugation of AMPs to larger antibody fragments is an elegant solution to overcome the shortcomings of peptide therapeutics like bioavailability, toxicity, and short half-life. Till date, antibiotics have been the most effective molecules for antibody drug conjugates, with rifamycin type antibiotics having been widely used (62). As we wanted to eliminate the complexity of linking the drug to the VHH by a separate chemical reaction that would increase the production costs considerably, we chose a linear peptide from humans, histatin, so that the AbTid could be produced as a fusion molecule in a single step from microbial production system. Histatin has been reported to neutralise Candida and all the ESKAPE pathogens equally well with different modes of action (26,27,28,30) but if used by itself is toxic to human cells. However, when fused with VHHs, the toxicity was eliminated as the larger antibody fragment was wrapped around it and the antimicrobial action was activated only when set free by the pathogen proteases. We used two versions of histatin, one full length and the other truncated form containing only the active site. Both the fragments worked equally well in neutralizing the pathogens. Hence a sort of design flexibility exists in the linker as well as the peptide, that can be exploited further to come up with a strategy for designing molecules with desirable physio-chemical properties like immunogenicity and solubility.

We characterized one of the AbTids - the anti-Pseudomonas ABG 14 further and found it to be resilient to adaptive resistance across nine generations. This was expected as it is a biological not susceptible to simple point-mutation based resistance due to its complex multiple modes of action – blockage of the metabolite transporter and bacteriolytic action of the AMP. The molecule did not destroy the beneficial bacteria that constitute a healthy patient microbiota due to its extreme specificity and also destroyed *P.aeruginosa* biofilms. It showed good *in vivo* efficacy and resulted in the survival of 75% of infected mice with corresponding decrease in the blood bacterial counts. All the three AbTids were at least 10 times more effective in neutralizing the pathogens compared to their corresponding VHHs, attesting to the mechanistic design efficiency making them suitable lead molecules for control of superbugs. Being a modular molecule, the peptide can be changed with a different linear peptide having better druggable properties and more tolerable toxicity profiles. New ‘hits’ can be generated in a matter of months by shuffling the VHH, peptide or linker and moved to lead optimization stage within a year, shortening the drug development process considerably.

## 5. CONCLUSION

In this study we demonstrate a novel non-traditional approach for developing new antimicrobials by using a combination of two biologicals: a pathogen specific neutralising camelid antibody fragment and an antimicrobial peptide. The antibody-drug conjugates are 10-20 times more effective than the antibody alone and neutralise pathogens irrespective of the drug resistance profiles with an efficacy in the sub-micromolar range. The modular nature and the complex mode of action make these molecules evolvable as well as resilient to simple point mutation-based resistance with an anticipated longer product life cycle. Being a camelid antibody fragment, the immunogenicity is expected to be low and the molecule can be humanised if required. These molecules exert their activity within 2-4 hours, a time during which it is expected to remain in circulation and concentrate at the site of action due to the binding activity of the antibody. Furthermore, the extreme specificity implies that AbTids will not disrupt the patient microbiota (reduced morbidity), ensuring activity only at site of action, resulting in faster recovery of patients in critical care settings. This class of molecule fulfils the three conditions for non-traditional therapeutics - specificity, evolvability and resilience to resistance (12). Although we have not demonstrated this concept against all the ESKAPE pathogens, but its similar efficacy in three different pathogens with different surface properties makes us believe that newer antimicrobials against the other pathogens as well as molecules with a broad-spectrum mode of action can be developed by choosing the appropriate targets and cleavage sites using this simple design principle. The VHH can be reformatted as well, and bivalent/bispecific molecules can be made quickly, considerably cutting the time and cost for developing a new molecule. Aided by a favorable regulatory environment, the strategy described here can be adopted to keep drug discovery research one step ahead of mutation-based resistance, which currently shows no signs of abating.

## 6. DECLARATIONS

### Ethics approvals and consent to participate

All applicable international, national, and/or institutional guidelines were followed. Experiments in AbGenics were carried out with Institutional Biosafety Committee Approvals.

### Consent for publication

Not applicable.

## Availability of data and materials

All data generated or analysed during this study are included in this article and its supplementary figures and can be provided by the corresponding author upon request.

## Authors contribution

SB formulated the concept and the therapeutic approach and oversaw the overall planning of the project. AN, PP and MB developed the protein purification strategies and executed the animal

## Acknowledgments

We are grateful to Dr. Vijay Satav of Dr. Satav’s pathological laboratories for providing us with the clinical samples of *P. aeruginosa* as well as the other pathogens for developing and testing the specificity of the camelid antibodies. Late Mr. Anil Nahar for constant encouragement and funding to build up the infrastructure required for the execution of the project. The authors acknowledge Mr. Siddharth Umarje for the critical evaluation of the manuscript, plotting the figures, statistical analyses of data and artistic depiction of the hypothesis, Mrs. Manisha Sabnis, and Mrs. Anamika Singh for the initial development of the phage display and protein purification protocols.

